# Double trouble: having two hosts reduces infection prevalence in vectored trypanosomatids

**DOI:** 10.1101/2021.04.29.440076

**Authors:** Al-Ghafli Hawra, M. Barribeau Seth

## Abstract

Trypanosomatids are a diverse family of protozoan parasites, some of which cause devastating human and livestock diseases. There are two distinct infection life-cycles in trypanosomatids; some species complete their entire life-cycle in a single host (monoxenous) while others infect two hosts (dixenous). Dixenous trypanosomatids are mostly vectored by insects, and the human trypanosomatid diseases are caused mainly by vectored parasites. While infection prevalence has been described for subsets of hosts and trypanosomatids, little is known about whether monoxenous and dixenous trypanosomatids differ in infection prevalence. Here, we use meta-analyses to synthesise all published evidence of trypanosomatid infection prevalence for the last two decades, encompassing 931 unique host-trypansomatid systems. In examining 581 studies that describe infection prevalence, we find, strikingly, that monoxenous species are two-fold more prevalent than dixenous species across all hosts. We also find that dixenous trypanosomatids have significantly lower infection prevalence in insects than their non-insect hosts. Within monoxenous trypanosomatids, genera infecting bees are most prevalent and infection prevalence does not vary between wild and managed bees. To our knowledge, these results reveal for the first time, a fundamental difference in infection prevalence according to host specificity where vectored species suffer from lower infection prevalence as a result of a ‘jack of all trades, master of none’ style trade-off between the vector and subsequent hosts.

## Introduction

Trypanosomatids are Kinetoplastid protozoan parasites that infect a diverse range of hosts including vertebrates, invertebrates and plants. Among trypanosomatids, there are two distinct life-histories, with some infecting only a single host (monoxenous), and others that require two different hosts to complete their development (dixenous). The dixenous lifestyle is thought to be evolutionarily derived and has evolved independently multiple times among the Kinetoplastids.^1^ Dixenous trypanosomatids, which are currently classified into five different genera (*i.e. Leishmania, Trypanosoma, Endotrypanum, Porcisia and Phytomonas*) invariably include an insect vector that then transmits infection to a vertebrate or plant host. The vast majority of epidemiological and empirical evidence for all trypanosomatids comes from the two medically relevant genera (*Leishmania and Trypansoma*). At least 20 species of *Leishmania,* cause cutaneous, mucocutaneous and visceral leishmaniasis; *Trypanosoma cruzi* causes Chagas disease and *Trypanosoma brucei* is the etiological agent of African sleeping sickness. These diseases cause more than 830,000 infections every year^2^ and their control is challenging. These parasites cause significant disease burden. Estimates of incidence and mortality vary for these diseases, but some studies predict that more than 1 million people contract leishmaniasis annually, including up to 700,000 visceral cases (with a fatality rate of 95%), and a further 1 billion people are at risk of infection.^3–5^

Monoxenous trypanosomatids not only have a simpler life-cycle and narrower host specificity, usually in a single invertebrate host, but they are also more common, more ancient (but see),^6^ and more diverse than dixenous trypanosomatids.^7^ There are 17 currently described monoxenous trypanosomatids genera: *Angomonas, Blastocrithidia, Blechomonas, Crithidia, Herpetomonas, Jaenimonas, Kentomonas, Lafontella, Leptomonas, Lotmaria, Novymonas, Paratrypanosoma, Rhynchoidomonas, Sergeia, Strigomonas, Wallacemonas,* and *Zelonia* in contrast to the four dixenous genera. Despite the overwhelming diversity of monoxenous trypanosomatids we know comparatively little about their natural history and prevalence, due to their perceived lack of medical relevance - although this has been called into question recently by opportunistic vertebrate infections by some monoxenous trypanosomatids.^8–10^

Monoxenous and dixenous trypanosomatids also appear to differ in their infection prevalence. Despite the substantive health burden of vectored trypanosomatids, they have surprisingly low infection prevalence in the vector insects. For instance, fewer than 3% of *Phlebotomus* sand flies carry *Leishmania*^11^ and as few as 0.1% of adult tsetse flies have transmissible *Trypanosoma* cells in their saliva.^12^ In contrast, trypanosomatids with only a single insect host can reach extremely high prevalence. In the bumblebee *Bombus terrestris* up to 35% of workers can be infected by its trypanosomatid parasite, *Crithidia bombi*.^13^ These data hint at something crucial: that perhaps the major limiting factor for trypanosomatid infection transmission is the vector, not the vertebrate host. Therefore, a detailed understanding of insect-parasite interactions, and how they can be manipulated, will be vital to the control and potential eradication of these diseases.

Here, we set out to systematically test whether monoxenous and dixenous trypanosomatids indeed have different infection prevalences, and whether this is consistent among hosts. While some systematic reviews have attempted to examine trypanosomatid prevalence they have only done so in the context of either individual trypanosomatid or host taxa,^14,15^ or alternatively summarised evidence across hosts but were geographically restricted.^16–18^ This is the first review of its type at this scale aimed to describe how fundamental aspects of parasite life-history influence disease prevalence in an important group of human, livestock, and wildlife parasites. We hope that our findings will help clarify the link between parasite life-history and infection in turn, potentially shed light onto better control strategies to combat the spread of trypanosomatid infections.

## Methodology

### Literature search

We searched Scopus and all available databases in Web of Science, to identify all published studies that describe trypanosomatid prevalence in non-human hosts (identifying 1,511 references in Scopus and 5,414 from Web of Science). Full search terms can be found in Supplementary Table 1. During the literature search, we did not apply any year or publication type restrictions (search stopped Jan 2020). To exhaustively find studies of neglected trypanosomatids, we further checked the references in relevant papers and added candidate citations to screening list of studies. All duplicate references were removed.

### Relevant screening stages and design

References were initially screened based on titles and abstracts manually by a single reviewer (HA) using Rayyan website as outlined in (Supplementary appendix)^19^. We further employed an automated second review protocol using Machine Learning Algorithm (MLA) and Natural Language Processing (NLP) during the initial screening phase, as has previously been suggested for large datasets^20^ (and detailed in (Supplementary appendix)). This was done to replace the second the reviewer before full-text screening. All studies that received a manual or an automated “include” decision by human reviewer or ML were subsequently subjected to further eligibility assessment by full-text review by two different reviewers (Figure.1). Inclusion and exclusion criteria, and detailed protocol of study selection and data extraction can be found in (Supplementary appendix).

**Figure 1:**
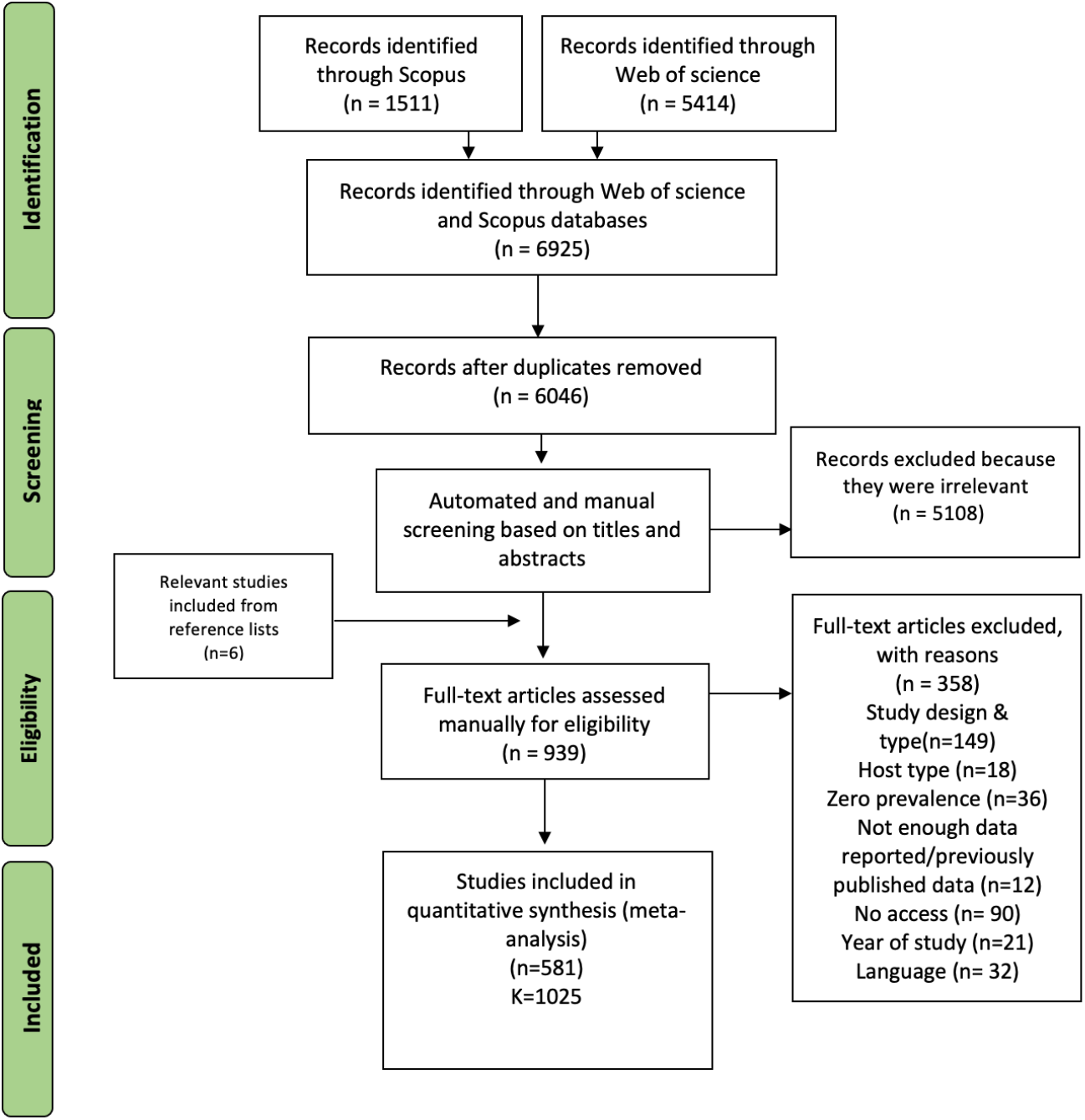
Selection and screening stages. The overall protocol utilised for literature search and screening, where (n) represents the number of studies included/excluded at each stage and (K) represents the number of data points extracted from included studies. All steps of the systematic review were in concordance with PRISMA guidelines.^22^

### Meta-analysis

We extracted the infection prevalence, diagnostic method, country of study, species of parasite and host, and year of publication from all included studies. As host species was not always provided, we simplified hosts to broad host categories (i.e. pigs, dogs, cats, camels, cattle, sheep, fish, raccoons, buffaloes, rodents, true bugs, flies, bees, birds, bats, fleas) when comparing across host groups. The proportion of infection values were first transformed with `escalc()` function and (measure=“PFT”) to stabilise the variance by Freeman-Tukey double arcsine transformation. Subsequently, meta-regressions were conducted on a trasformed scale using “metafor” package. Pooled estimates (weighted proportions) were then back-transformed for each analysis and multiplied by 100 to get the pooled prevalence. The pooled prevalence can be interpreted as weighted summary of infection prevalence for that group. Publication bias was assessed using a funnel plot with trim and fill method and Egger’s test with sample size as a predictor. We employed a random effect multilevel model (rma.mv) with a nested random design to account for the heterogeneity among studies (e.g. study protocols and/or different host/parasite taxa within and between studies). We assigned “study ID” as an outer random factor, and both host type (19 levels) and parasite taxa (50 levels) were considered inner-random factors. Each model was fitted with restricted maximum-likelihood (“REML”). This was done for the general model with no moderator and meta-regressions with the following moderators: diagnostic method (3 levels [“molecular”, “microscopic & culture based”, “serological”]), life-history traits of the parasite (3 levels [dixenous, monoxenous, mixed trypanosomatids), host groups (insects and non-insects), parasite taxa (50 levels), host taxa (19 levels). We also ran similar meta-regression models to assess pooled prevalence on subgroups of interest (as detailed in Supplementary Table 2).

Between-study variation was assessed using the Higgins 12 statistic using (“rma” or “rma.uni”), which usually indicates substantial heterogeneity if exceeding 50%. The QM statistic, an omnibus test of all model coefficients except for the intercept, was calculated to conclude if a moderator significantly influences the mean effect size. All meta-regression models were run with and without the model intercept to statistically compare effect size of factors and generate pooled estimate for each factor respectively. We used a stringent Benjamini-Hochberg method to account for elevated Type 1 error from multiple testings (Supplementary Table 3). Forest plots were presented using “orchaRd” package with “tanh” transformation as traditional forest plot will fail to present the large number of studies used here.^21^ Data sources for each analysis is outlined in (Supplementary Appendix).

We used several distinct meta-regressions (presented in Supplemental table 2) to address particular questions. In most models, we ran the meta-regression both without, and with an intercept allowing us to compare levels of the moderator using t-tests. Each metaregression asks a specific question of these data. In short, model A examined variation in infection prevalence among parasite taxa, model B asks whether parasites vary in infection prevalence depending on whether they infect a single host or two hosts, model C compares infection prevalence in insects to non-insect hosts. The remaining models further delve into differences within host or parasite taxa (Supplementary Table 2). Models F1 and F2 examine variation in diagnostic methods in insect (F1) or non-insect (F2) hosts; F3-F5 compare dixenous to monoxenous trypanosomatids in all insects, flies, or true bugs respectively; F6-F9 compare prevalence of insect hosts to non-insect hosts in different levels of dixenous trypanosomatids (all dixenous trypanosomatids F6, Leishmania spp. F7, Trypanosoma spp. excluding T. cruzi F8, T. cruzi only F9) and F10 - F12 examine wild vs managed bees combined, bumblebees, and honeybees, respectively. All models included study ID as an outer random factor while the inner random factors varied (A-F5: host type and parasite type; F6-9: parasite taxa; F10-F12: bee taxa and parasite taxa). All models (except A) included an intercept, which allows us to statistically compare between inner factors of the moderator with t-test. We did not include an intercept in meta-regression A as we were less concerned about how much each level differed, but rather the overall effect of a moderator on infection prevalence. Further details about the meta-regressions can be found in the supplemental methods and supplemental table 2.

### Results

From our initial input of 6046 unique citations, we retained 581 studies after full-text screening. In general, these studies demonstrate a wide geographic distribution of trypanosomatids (Supplementary Appendix) among a diverse range of host-taxa (Fig.2). For summary purposes and to capitalize on this large data-set, we show a summary of infection prevalence among different parasite-host systems in (Supplementary Fig. 2), subsequent analysis of pooled estimate of various host-groupings in (Supplementary Fig.7), infection prevalence stratified by parasite species in(Table 1), or by host type in (Supplementary Fig.4). Some host groups (e.g. fish, raccoons and buffaloes) have a very high pooled prevalence although they are comparatively rarely studied (Supplementary Fig.4). In spite of the low number of studies, their total sample size indicates comparable robustness of analysis to that of the otherwise heavily examined host groups for trypansomatid infections (Supplementary Fig.7).

**Figure 2:**
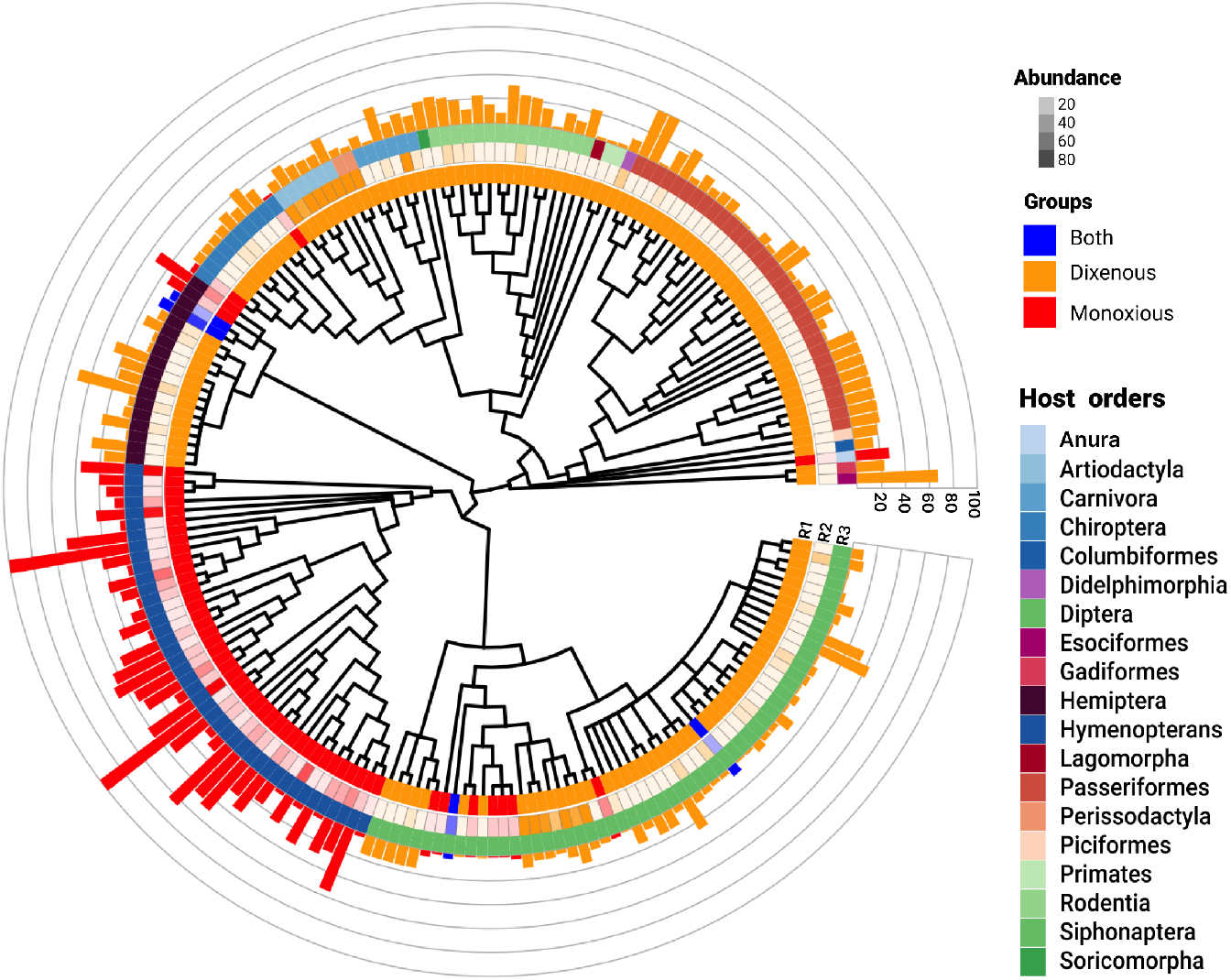
A visual summary of all included data among various host-trypanosomatid systems. A phylogenetic representation of all host taxa examined in this study. The first (inner) ring (R1) illustrates the parasites’ life-history (dixenous, monoxenous or both), the second ring (R2) highlights the abundance of prevalence data for each taxon, and the third ring (R3) is host order. Prevalence of infection (%) is shown in the outer scaled ring, which is coloured according to parasite life-history.

**Table 1.**
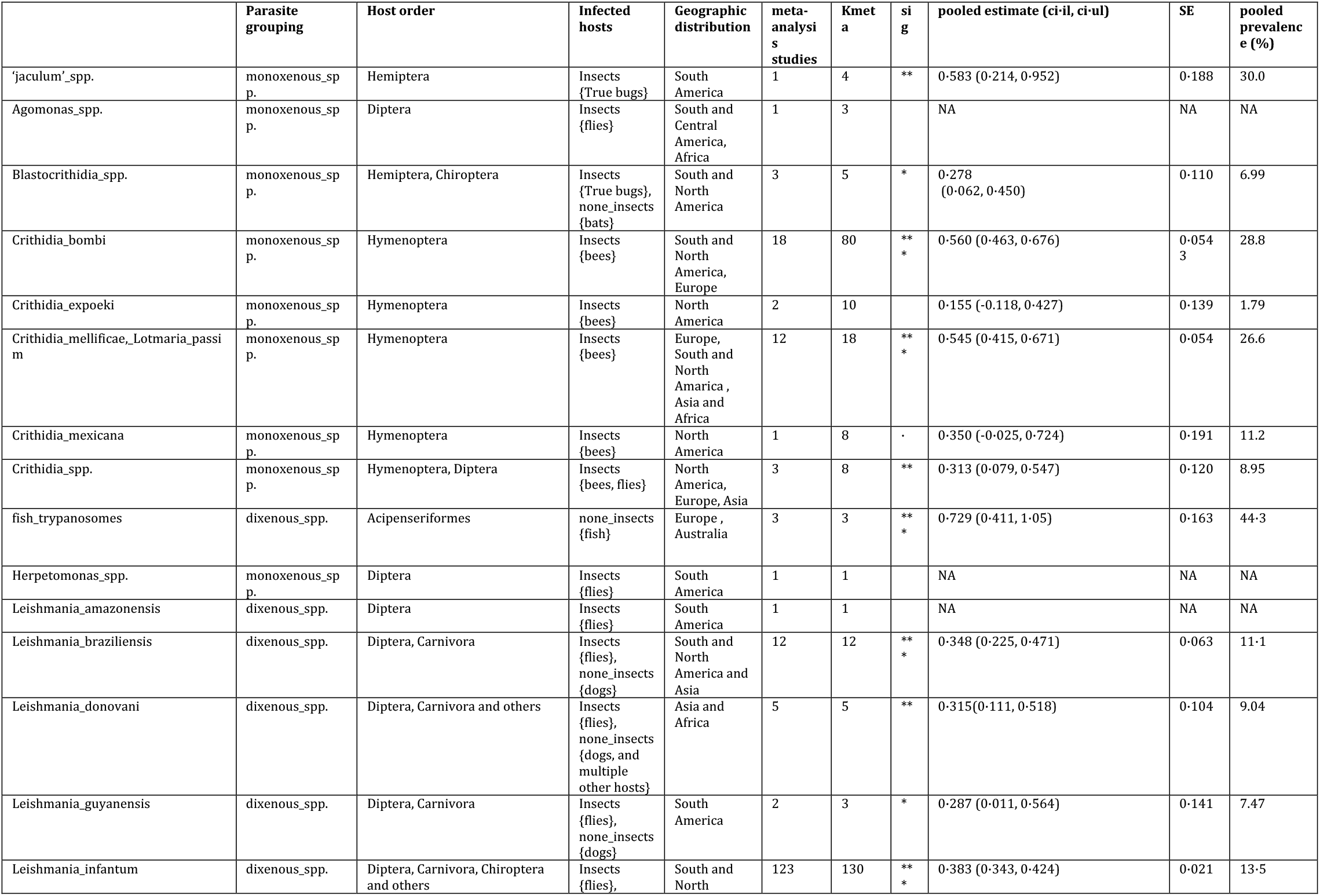
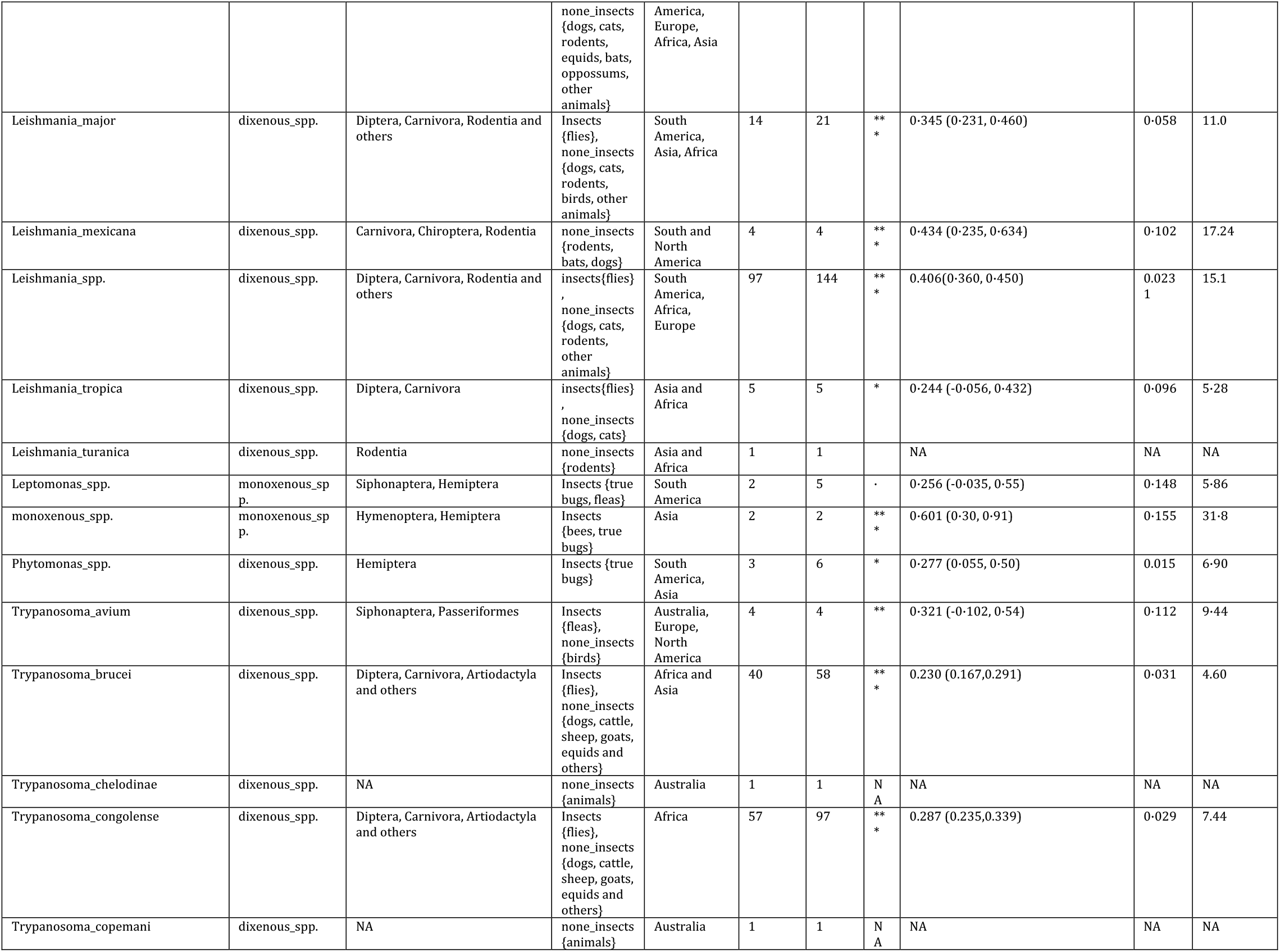
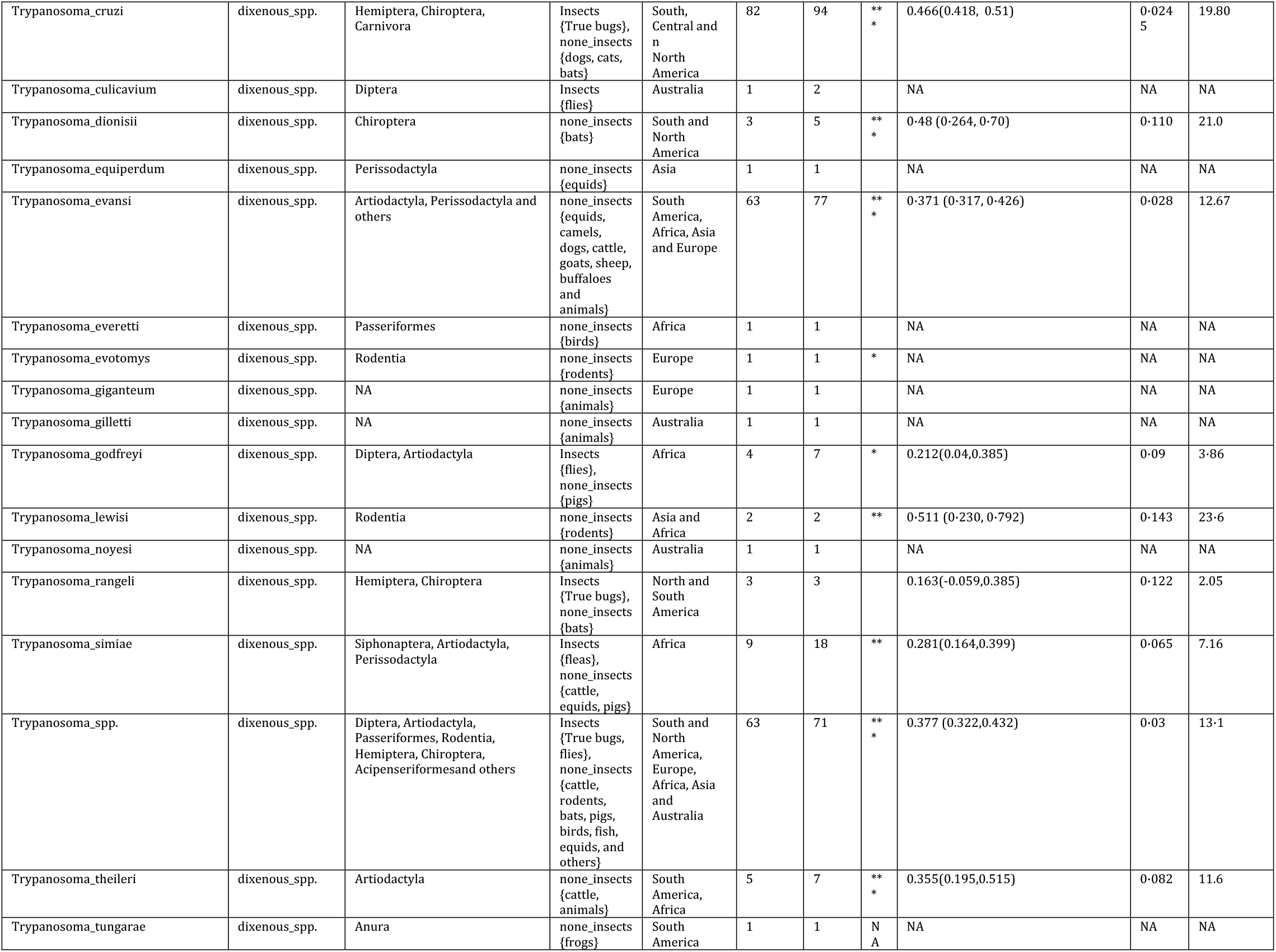
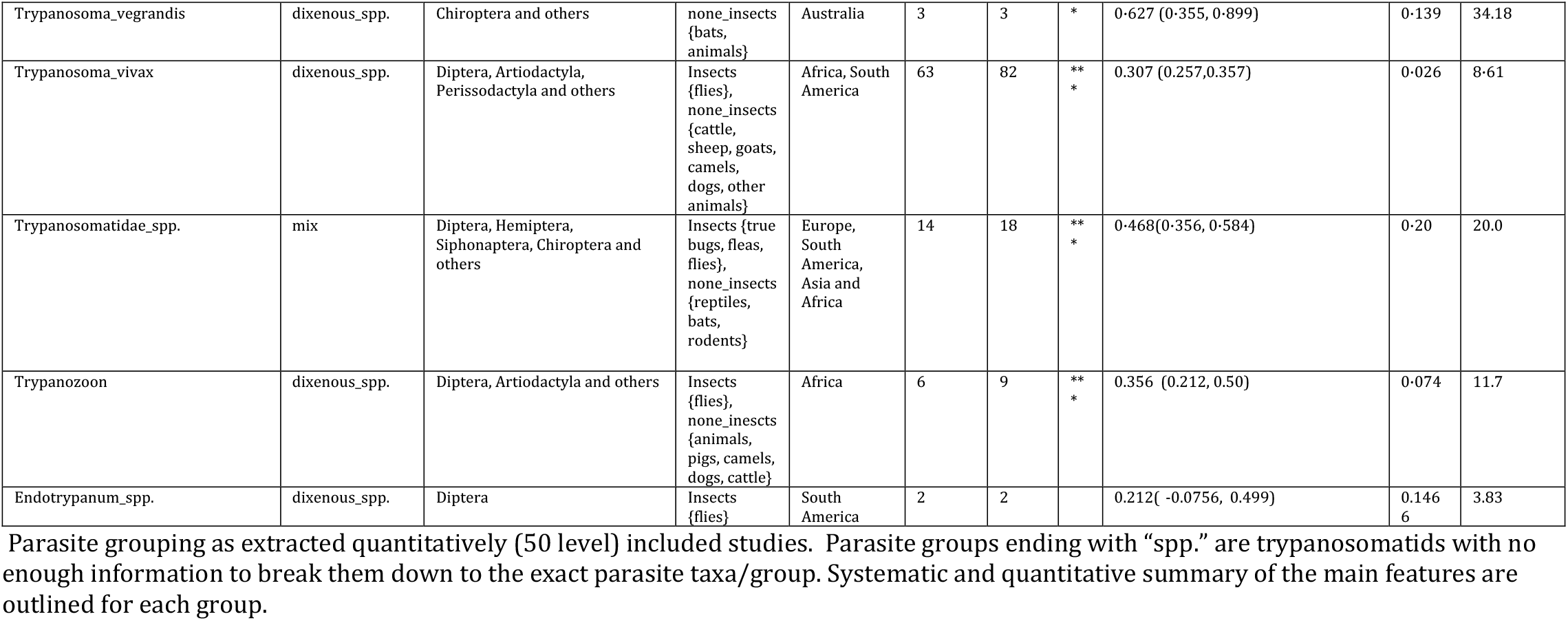
Summary of all included studies based on parasite groupings.

[monoVdix] One of the main aims of this work was to quantitatively examine, for the first time, the impact of life-history traits on infection prevalence. Thus, we sat to test whether pooled prevalence of monoxenous infections differ from dixenous infections (both among insect and non-insect hosts). We found a significant difference (*P* = 0.0067) between pooled prevalence of monoxenous (20.6%) and dixenous infections (13.2%) (Fig.3A). Introducing “host-traits” as a sole moderator (i.e. insects and non-insects) reveals no significant difference (after correcting for multiple testing) between pooled prevalence of the two groups, 15% and 13.3% respectively (Fig.3B). Limiting the analysis to insects alone revealed similar trend where monoxenous infections are approximately two-fold more prevalent (21.9%) than dixenous infections (11.1%) (Fig.4, Table 2). We then compared prevalence in the groups of insects that host both dixenous and monoxenous parasites (true bugs and flies). Here however, we found non-significantly higher pooled prevalence for dixenous infections in flies and similar pooled prevalence in true bugs (Table 2).

**Figure 3:**
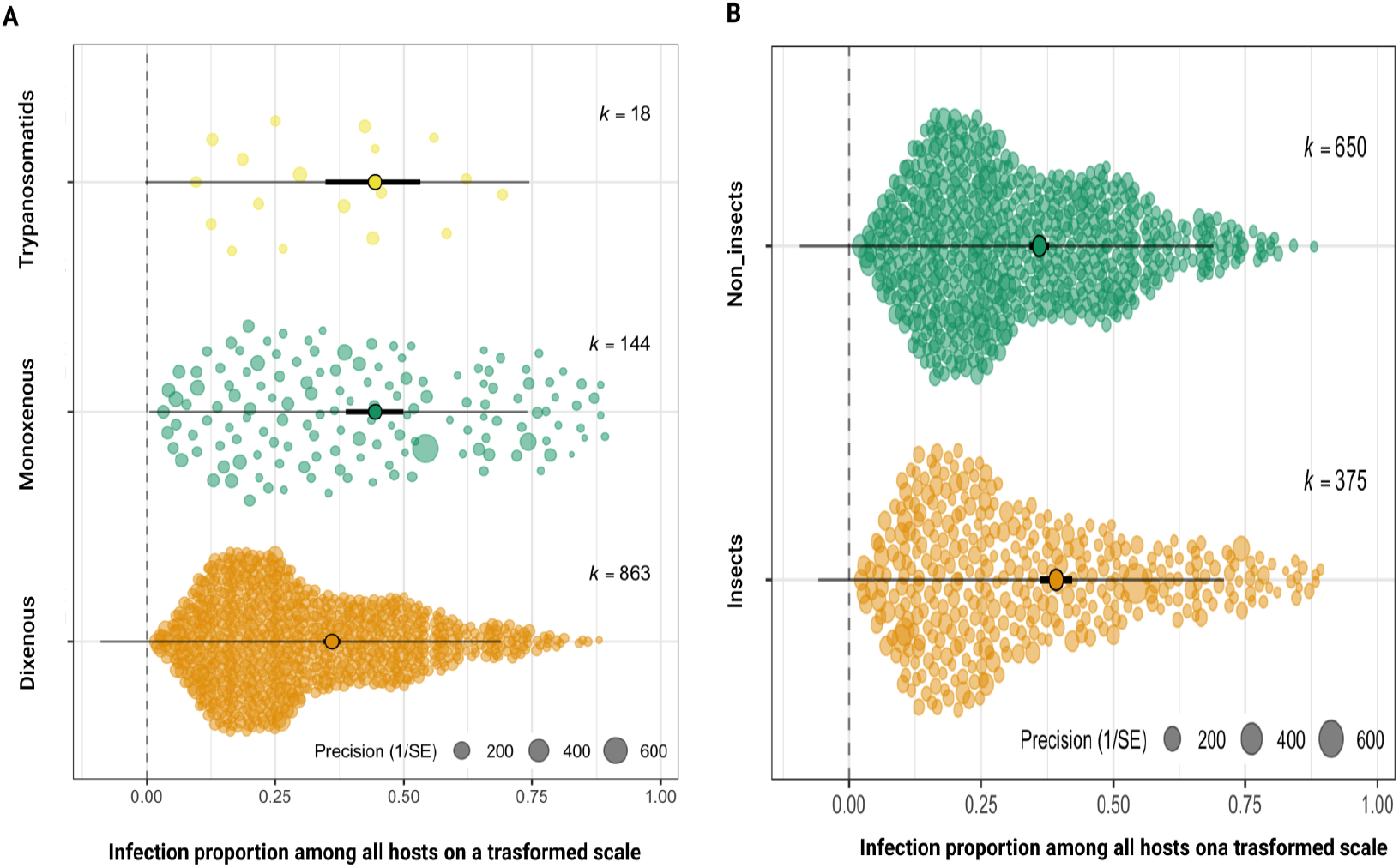
Orchard plots of infection prevalence among trypanosomatids that vary according to the number of hosts they infect (A) and whether those hosts are insects or non-insects (B). The X-axis shows effect sizes for each study (circles). Random jitter was added on the Y-axis to aid visibility. Pooled effect size (proportion of infection) is shown in the middle accompanied with bold lines representing the 95% CI. A) Pooled estimate of infection prevalence of dixenous (0.377, 95% CI:0.358, 0.40), monoxenous (0.478, 95% CI:0.409, 0.548), and mixed trypanosomatids (0.478, 95% CI:0.363, 0.593) with significant difference between dixenous and monoxenous (*P* = 0.0057, Benjamini-Hochberg adjusted cut-off *P* = 0.0188). These pooled estimates back-transform to 13.2% (dixenous), 20.6% (monoxenous), and 18.6% (mixed trypanosomatids). B) There is no overall significant difference between the pooled estimate of infection prevalence of insect (0.414, 95% CI: 0.378, 0.450) and non-insect (0.377, 95% CI:0.356, 0.398) hosts (*P* > 0.016;Benjamini-Hochberg adjusted cut-off). These pooled estimates correspond to 15.0% and 13.3% infection prevalence. These pooled estimates correspond to 15.0% and 13.3% infection prevalence.

**Figure 4:**
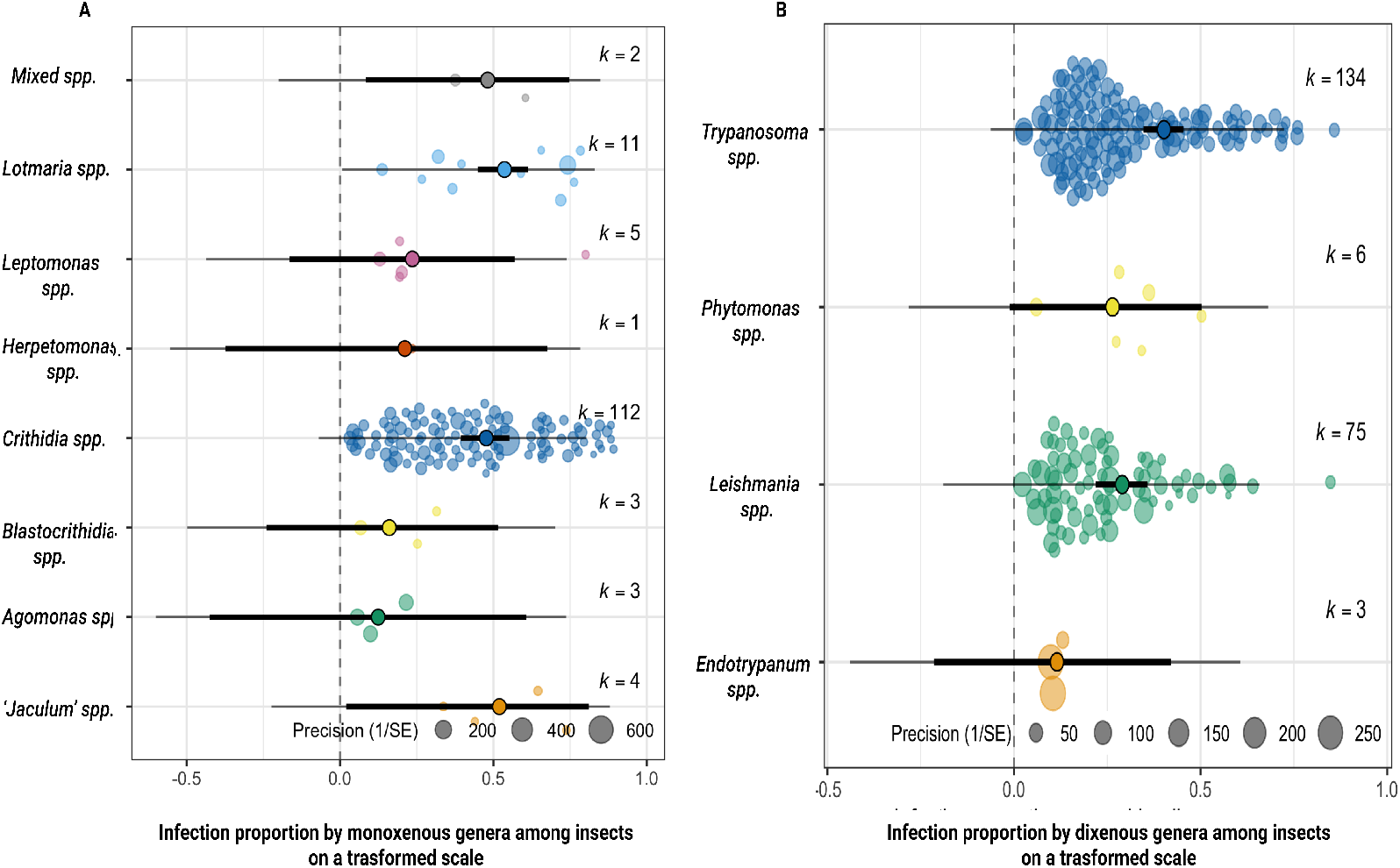
Orchard plots of monoxenous (A) and dixenous (B) parasites prevalence in insect hosts. The X-axis shows effect sizes for each study (circles). Random jitter was added on the Y-axis to aid visibility. Pooled effect size (proportion of infection) is shown in the middle accompanied with bold lines representing the 95% CI. A) Pooled estimate of infection prevalence of monoxenous genera, representing a pooled estimate of 0.525 (CI:-0.084, 0.965), 0.599 (CI:0.484, 0.713), 0.240 (CI:-0.167, 0.647), 0.215 (CI:-0.392, 0.821), 0.519 (CI:0.417, 0.621), 0.162 (−0.246, 0.570), 0.125 (CI: −0.455, 0.705) and 0.576 (CI:0.0199, 1.132) for the monoxenous species, *Lotmaria, Leptomonas, Herpetomonas, Crithidia, Blastocrithidia, Angomonas and ‘jaculum’* species respectively. The infection prevalence did not differ significantly among monoxenous species of trypanosomatids. These pooled estimates correspond to a prevalence of 24.1%, 31.0%, 3.94%, 2.82%, 23.5%, 0.994%, 0.185% and 28.8%. B) Pooled estimates of infection prevalence in dixenous genera, of 0.116 (CI:0.217, 0.449), 0.298 (CI:0.223, 0.374), 0.270 (CI:-0.0121, 0.553) and 0.426 (CI:0.362, 0.490) for *Endotrypanum, Leishmania, Phytomonas* and *Trypanosoma* corresponding to prevalence of 0.948%, 8.30%, 6.78% and 16.8%. The infection prevalence did not differ significantly among dixenous species of trypanosomatids (*P* > adjusted cut-off based on Benjamini-Hochberg correction).

**Table 2:**
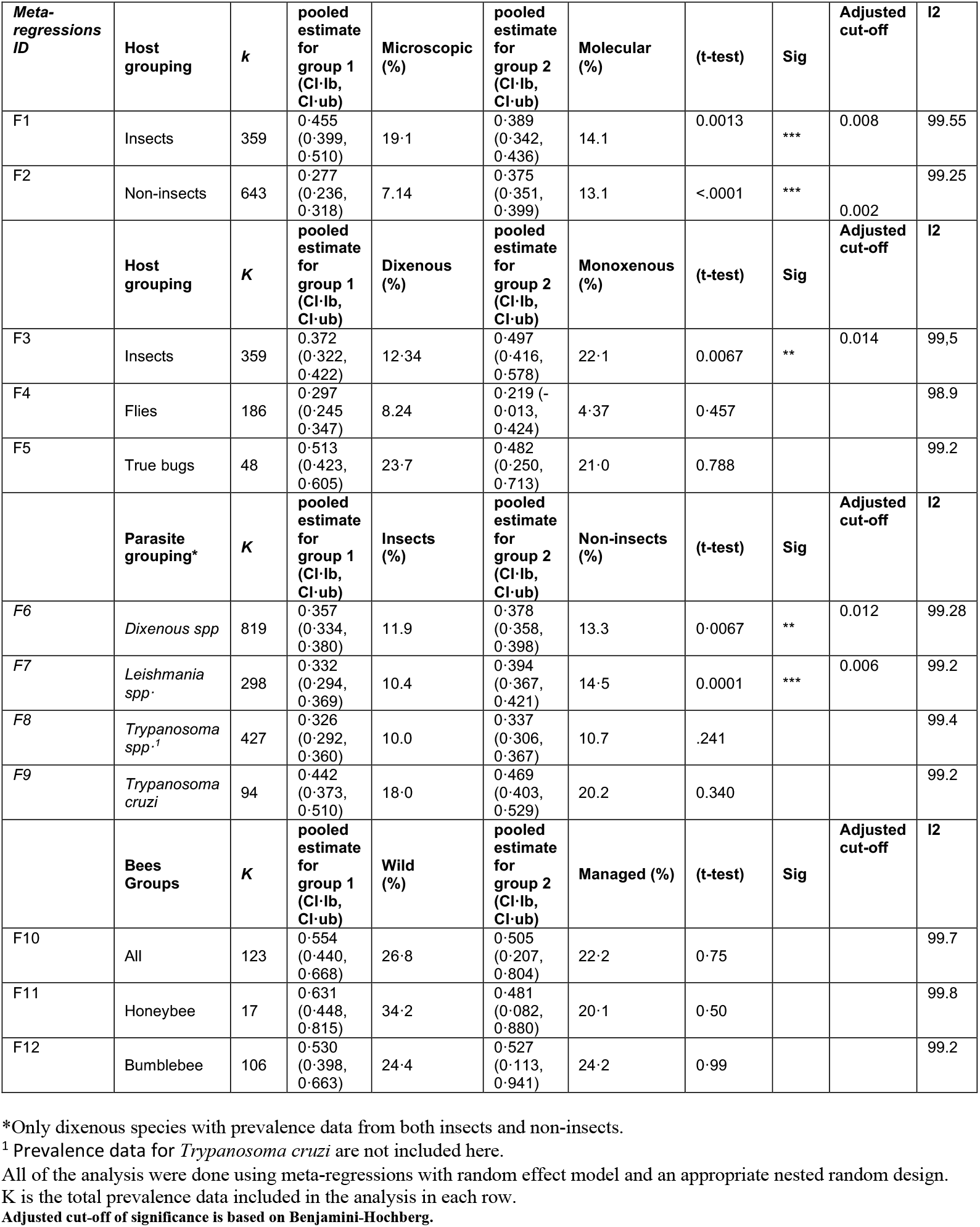
Meta-regressions on subgroups of interest.

| **Meta-regressions ID** | **Host grouping** | **k** | **pooled estimate for group 1 (CI-lb, CI-ub)** | **Microscopic (%)** | **pooled estimate for group 2 (CI-lb, CI-ub)** | **Molecular (%)** | **(t-test)** | **Sig** | **Adjusted cut-off** | **I2** |
| --- | --- | --- | --- | --- | --- | --- | --- | --- | --- | --- |
| F1 | Insects | 359 | 0.455<br>(0.399, 0.510) | 19.1 | 0.389<br>(0.342, 0.436) | 14.1 | 0.0013 | *** | 0.008 | 99.55 |
| F2 | Non-insects | 643 | 0.277<br>(0.236, 0.318) | 7.14 | 0.375<br>(0.351, 0.399) | 13.1 | <.0001 | *** | 0.002 | 99.25 |
|  | **Host grouping** | **K** | **pooled estimate for group 1 (CI-lb, CI-ub)** | **Dixenous (%)** | **pooled estimate for group 2 (CI-lb, CI-ub)** | **Monoxenous (%)** | **(t-test)** | **Sig** | **Adjusted cut-off** | **I2** |
| F3 | Insects | 359 | 0.372<br>(0.322, 0.422) | 12.34 | 0.497<br>(0.416, 0.578) | 22.1 | 0.0067 | ** | 0.014 | 99.5 |
| F4 | Flies | 186 | 0.297<br>(0.245, 0.347) | 8.24 | 0.219 (-<br>0.013, 0.424) | 4.37 | 0.457 |  |  | 98.9 |
| F5 | True bugs | 48 | 0.513<br>(0.423, 0.605) | 23.7 | 0.482<br>(0.250, 0.713) | 21.0 | 0.788 |  |  | 99.2 |
|  | **Parasite grouping\*** | **K** | **pooled estimate for group 1 (CI-lb, CI-ub)** | **Insects (%)** | **pooled estimate for group 2 (CI-lb, CI-ub)** | **Non-insects (%)** | **(t-test)** | **Sig** | **Adjusted cut-off** | **I2** |
| F6 | *Dixenous spp* | 819 | 0.357<br>(0.334, 0.380) | 11.9 | 0.378<br>(0.358, 0.398) | 13.3 | 0.0067 | ** | 0.012 | 99.28 |
| F7 | *Leishmania spp.* | 298 | 0.332<br>(0.294, 0.369) | 10.4 | 0.394<br>(0.367, 0.421) | 14.5 | 0.0001 | *** | 0.006 | 99.2 |
| F8 | *Trypanosoma spp.*<sup>1</sup> | 427 | 0.326<br>(0.292, 0.360) | 10.0 | 0.337<br>(0.306, 0.367) | 10.7 | .241 |  |  | 99.4 |
| F9 | *Trypanosoma cruzi* | 94 | 0.442<br>(0.373, 0.510) | 18.0 | 0.469<br>(0.403, 0.529) | 20.2 | 0.340 |  |  | 99.2 |
|  | **Bees Groups** | **K** | **pooled estimate for group 1 (CI-lb, CI-ub)** | **Wild (%)** | **pooled estimate for group 2 (CI-lb, CI-ub)** | **Managed (%)** | **(t-test)** | **Sig** | **Adjusted cut-off** | **I2** |
| F10 | All | 123 | 0.554<br>(0.440, 0.668) | 26.8 | 0.505<br>(0.207, 0.804) | 22.2 | 0.75 |  |  | 99.7 |
| F11 | Honeybee | 17 | 0.631<br>(0.448, 0.815) | 34.2 | 0.481<br>(0.082, 0.880) | 20.1 | 0.50 |  |  | 99.8 |
| F12 | Bumblebee | 106 | 0.530<br>(0.398, 0.663) | 24.4 | 0.527<br>(0.113, 0.941) | 24.2 | 0.99 |  |  | 99.2 |

\*Only dixenous species with prevalence data from both insects and non-insects.
<sup>1</sup> Prevalence data for *Trypanosoma cruzi* are not included here.
All of the analysis were done using meta-regressions with random effect model and an appropriate nested random design.
K is the total prevalence data included in the analysis in each row.
Adjusted cut-off of significance is based on Benjamini-Hochberg.

Notably, dixenous parasites infect 1-17% of their insect hosts with no significant difference among parasite genera (Fig.4B). Common dixenous parasites, with prevalence data from both insects and non-insects, were slightly but significantly more prevalent in their noninsect hosts (13.3%) compared to their insect hosts (11.9%) (Table 2).

Monoxenous genera exhibit two alternative trends: high pooled prevalence for *Lotmaria, jaculum* and *Crithidia* species, but low pooled prevalence for *Herpetomonas, Leptomonas, Agomonas,* and *Blastocrithidia* groups (Fig.4B). Taking into consideration that a considerable amount of monoxenous data (80%) were on bee infections, we performed an meta-regression on subgroups to further understand the main drivers of bee infection prevalence. While the majority of included data represent natural infection among wild bees, the remaining evidence describes spillover of infections e.g. by sampling non-native/commercial bees. We did not find any difference in infection prevalence between wild-bees and commercial/managed bees for bumblebees or honeybees (Table 2).

Honeybee infections seem to predominantly occur in Western/European honeybees (i.e. *Apis mellifera Linnaeus*) compared to Asian honeybees (e.g. *Apis dorsata Fabricius* and *Apis cerana Fabricius*). We also noted a significant reduction in heterogeneity of evidence across all honeybee-infections due to spatial variation only (46% heterogeneity). However, a much higher diversity of infection prevalence was found in bumblebee taxa. Accounting for spatial and temporal factors failed to significantly reduce the heterogeneity in evidence for bumblebee infections, suggesting a unique impact of various bumblebee-trypanosomatids systems on different patterns of prevalence (Fig.3.B) and likely the greater diversity of bumblebees.

Diagnostic approach also affected infection prevalence but differently according to host. Microscopic diagnostics revealed higher infection prevalence in insects whereas molecular assays report higher infection prevalence in non-insects (Table 2).

## Discussion

Trypanosomatid parasites have two distinct life-histories that vary in the number of host species necessary to complete their life-cycle. This difference is profound and will have commensurately profound effects on the evolution of parasite traits, including their ability to establish infection. How these alternative life-history strategies influence infection prevalence, however, is poorly understood. To our knowledge, this is the first comprehensive study designed to test for differences in infection prevalence.

We found that monoxenous trypanosomatids are consistently more prevalent, by two-fold, than dixenous species in insects alone (Table 2) and similarly across all hosts (Fig. 3A). Taken together, one cannot escape the impression that complex parasite life-cycles have a cost to infection prevalence. In line with such speculation, high prevalence for monoxenous species was mainly notable among genera that infects bees such as *Crithidia* and *Lotmaria* with pooled prevalence of 23.9% and 24.9% respectively; ranking bees with the highest pooled prevalence among all insects (27.0%). It is striking to realize that about a quarter of surveyed bees are infected by trypanosomatid pathogens. This is of a particular concern considering the crucial role of bees as pollinators, which, with other insect pollinators, account for the fertilization of 83% of flowering plants^23^ including nutritionally and economically important crop species.^24,25^ Bees and other pollinators have been declining. Among the presumed causes of pollinator declines are infectious diseases.^26^ Despite the attention that bee-trypanosomatid infection has attracted, the distribution of interest remains uneven. Infection prevalence of trypanosomatids is largely unexplored outside of Europe and North America (Supplementary Fig.5)

The relationship between wild and managed bees, and the risk of spillover in either direction, has received particular attention and concern in the literature. While managed bees have been blamed for parasite transmission to wild bees,^27,28^ we found no significant difference between managed and wild honeybees and bumblebees (either independently or combined) in infection prevalence. Whether this reflects shared susceptibility to infection or ongoing spillover to and from wild pollinators remains unknown.

To our knowledge this is the first study to provide a full summary of prevalence data among all host-trypanosomatids systems. Encouragingly, our findings recapitulate other taxon- or geographically-restricted systematic reviews. For instance, a similar range of prevalence (2.4-9.2%) for trypanosomatids was reported in sand-flies (*Phlebotomus* spp.) in Iran,^29^ which agrees with our finding of 10.14% (Table 2, meta-regression F7). Another meta-analysis on trypanosomatids in tsetse flies revealed a prevalence of 10%, as did we in flies infected by *Trypanosoma*.^15^ Thus, although biting flies are responsible for most human trypanosomatids, our results seem to comport with previous studies revealing low infection prevalence in flies as a group, with no significant difference between monoxenous and dixenous parasites within flies (Table 2). Meanwhile, pooled prevalence for trypanosomosis in cattle, sheep and goats was reported as ranging between 1-9%, which is comparable to what we find (meta-regression F8 in Table 2, and Supplementary Fig.7).^17,18,30^

We also reveal significantly higher prevalence in the vertebrate hosts than vector flies, consistently across genera (Table 2). This brings to the fore a possibility that dixenous trypanosomatids infect more of their vertebrate hosts than their insects vectors. Evolutionary theories predict a life history trade-off between investment in survival (within-host persistence and growth) and reproduction (between-host transmission). For *Trypanosoma brucei* and other dixenous trypanosomatids; such relationship is amplified by the number of host species needed to complete the life-cycle.^31^ This trend warrants further investigations as understanding how these species are maintained in the multi-host metapopulation holds great potential in managing the spread of vector-borne diseases. For instance, if the prevalence of infection in vectors is low, further reduction in insect hosts may better control infection in the metapopulation than suppression in vertebrate hosts.

In this study, we found that microscopic and culture based diagnostics revealed higher infection prevalence in insects but greater infection prevalence was revealed from molecular assays in non-insects (Table 2). This result does not necessarily indicate greater accuracy for either assay, but may rather indicate the preferred methodology of researchers working with insect versus vertebrate hosts. In line with this explanation, Abdi and researchers^15^ concluded that microscopic diagnostics were more common in surveys of tsetse flies than molecular assays and Aregawi *et al.*^14^ reported a higher range of prevalence (6-28%) by molecular tests in vertebrates compared to microscopic tests (2-9%) for the detection of *Trypanosoma evansi.* Standardising the diagnostic method applied for the detection of trypanosomatids within insects and non-insects is crucial consideration for future research to enable accurate comparisons of prevalence that is unlikely due variations in diagnostic protocols. We took this factor into consideration while doing all meta-regressions (either on subgroups or the full dataset) presented in this study to ensure that the difference in prevalence accounts for variable diagnostics. To our assurance, in all of the main subgroup analysis, there was no significant difference in pooled prevalence due to different diagnostic tools when examining dixenous trypanosomatids among insects and non-insects, and individual groups of vertebrate hosts (e.g. cats, dogs, rodents, camels, goats, pigs, bats and buffaloes).

The key takeaway from this study is the unique pattern of infection prevalence in association with different life-history traits of trypanosomatids: revealing both (I) higher pooled prevalence for monoxenous species in insects and across all hosts, and (II) consistently significant lower prevalence among vectors compared to subsequent hosts for dixenous species. We present three non-mutually exclusive arguments that could explain this finding:

- First, monoxenous species are subject to selection pressure from only one host, thereby affording these species more opportunity to adapt to this host than parasites that must co-evolve with both an insect vector and an additional, often vertebrate, host. We would thus expect to see greater parasite fitness in specialised monoxenous parasites than in dixenous parasites as the monoxenous parasites would arrive closer to their evolutionary optima. Dixenous parasites, in contrast, may be forced to trade-off infection relevant traits in one host for traits in the second host.
- Second, these finding might be due to an equal force of infection shared among differing numbers of host species. Here, we offer a numerical explanation, where the same hypothetical number of infectious units are simply shared among more host species in dixenous parasites, thus diluting the infection prevalence in any one host species - to approximately half. This explanation is more than passingly similar to the “dilution effect” which is often, and sometimes controversially, used to describe infection risk in human populations when the natural host is not usually the human - e.g. Lyme disease,^32–34^ although to our knowledge it has not been invoked to explain variation in infection prevalence among vectored and non-vectored parasites.
- Finally, host characteristics such as lifespan, recruitment rate, sociality, and physiology may explain some of the variation in infection prevalence. For instance, social bees, which have higher infection prevalence, differ from the rest of insect hosts not only in their sociality but also in their lifespan. The biological unit for social bees is the colony, which can extend the lifespan of an infection to at least a year (bumblebees) and to several years in honeybees. The other insect hosts live for up to a few months, and have high birth and death rates which, along with rarer direct interactions can suppress the overall prevalence of infection regardless of the parasite life-cycle strategy. Consistent with this view, we found low pooled prevalence among flies with no significant difference between monoxenous and dixenous species. Furthermore, flies might be physiologically harder to infect by trypanosomatids than bees as genomic studies have described a reduced immune repertoire in bees relative to dipterans.^35,36^

Dixenous parasitism in trypanosomatids had evolved multiple times and million of years ago.These lineages have clearly been successful, persevering for an estimated 100 million years in the Leishmaniinae.^37,38^ Thus, despite the disadvantage imposed by the complexity of life-cycle that we describe here (i.e. reduced infection prevalence compared to their monoxenous kins), there must be other benefits to the dixenous life-history that selected for the transition to and the maintenance of this complex life-cycle. The tautological benefit of a dixenous life-cycle, is having two hosts. Having multiple hosts allows several potential benefits. The greater division between investment in growth (e.g. vertebrate host) and sexual reproduction (largely in insect vector host)^39,40^ may optimize the processes of both. This, for instance, allows greater clonal amplification in resource-rich large-bodied hosts and uses abundant highly mobile vectors for transmission. Maintaining infection in a long-lived host may also buffer parasites from the boom-and-bust population dynamics, seasonality, and high extrinsic mortality rates of insect hosts. Finally, transmitting to many vectors may allow occasional co-infection with other genotypes of parasite, and thus outcrossing potential.

It is worth noting that variation in transmission route has been implicated in other key parasite traits, most notably, in virulence. Ewald’s landmark paper^41^ proposed that vectored parasites are freed from the tradeoff between transmission and virulence. Support for this attractively intuitive hypothesis has been both theoretically and empirically mixed.^42–46^ While we did not examine virulence here, we were able to avoid one criticism of Ewald’s original work^47^ by comparing only among closely related parasites and including both host and parasite as random effects. We found distinct differences in infection prevalence depending on transmission route, demonstrating that broad scale parasite features, such as prevalence, or virulence, can be detected with such approaches.

While this study was ambitious in scope and exhaustive in design, there are limitations. Foremost is the small number of studies describing prevalence in monoxenous species. This is likely due to scarcity of publications for neglected trypanosomatids rather than study selection/handling approaches employed here. This study included all English published reports of trypanosomtids systems from the last 20 years that contribute to slyvatic transmission cycles, except studies that only examined human medical infections. Despite the exclusion of human infections, their impact and contribution to the transmission cycles either directly (to insects) or indirectly (to non-insects) have been accounted for by including studies surveying animals living nearby to humans. This includes pets (dogs, cats), domesticated animals, and the vectors, such as true bugs (tritomines), which are particularly found in houses. Thus, although the direct interaction in humans-trypanosomatids systems falls beyond the scope of this study, what we show here (i.e. how trypanosomatids differ in their infection prevalence among various nonhuman hosts) is particularly important to enable effective prevention plans for neglected tropical diseases of humans and livestock. Understanding the fundamental processes that determine infection prevalence will help us better understand trypanosomatids in general and identifying “weak-links” in the infection chain of vectored parasites - here in the vectored species, will enable better tailored control strategies for trypanosomatid diseases relevant to human, livestock, and wildlife health as well as food-security.

