## Supplementary Appendix for "Double trouble: having two hosts reduces infection prevalence in vectored trypanosomatids"

### Supplementary material

#### Extended protocol

##### Inclusion and exclusion criteria

A publication was provisionally included if it (I) was published in English between 2000 to 2020 with cross-sectional or prospective (longitudinal) design, (II) contains data on any positive diagnostic test for trypanosomatids in any naturally infected non-human host, (III) contains all necessary information to assess prevalence or enough information to calculate prevalence independently, (IV) describes the diagnostic test used.

Conversely, a publication was excluded if it failed to meet the above criteria or reports data derived from secondary sources (e.g. publicly available databases or previously published data), (I) reports data on human infections or based on experimental infections only, (II) is not an original paper (e.g. reviews and letter to editors), (III) is based on clinical signs only with no diagnostic tests conducted to confirm prevalence, (IV) has insufficient information to calculate the prevalence independently or reports zero infection by a diagnostic test for trypanosomatids.

All hosts infected by single or multiple species were included, similarly, species infecting different hosts or single hosts were also included. Prevalence data was broken down to the taxa level of both the host and the parasite whenever its possible based on reported information.

##### Manual screening based on titles and abstracts

A total of 6046 studies were entered to Rayyan website to initiate the manual screening process.[1] Reviews, book-chapters, letter to editors, single case studies and papers reporting only human cases or experimental infections have been identified using a list of relevant keywords/phrases (e.g "experiment", "experimentally infected", "retrospective", "review", "meta-analysis", "systematic review", "in vitro", "in vivo", "drug", "evolution", "treatment", "clinical trials", "human", "patients", "individuals", etc.). This generated a list of potentially irrelevant studies to be screened manually based on titles and abstract. A similar approach was employed to guide the initial screening of potentially relevant and eligible citations using keywords/phrases that include but not limited to the following: "survey", "cross-sectional", "prospective", "natural infection", "prevalence", "occurrence" and "field study". All studies then were subjected to full manual screening based on titles and abstracts, and no decision was made based on keyword calling only.

##### Automated screening based on titles and abstracts

The primary dataset of citations, encompassing 6046 studies, was further processed before MLA-screening. This was done by removing all publications that would not necessarily be excluded by NLP and MLA alone, such as due to irrelevant year of publication, duplication and wrong study type (e.g reviews, single case studies, and letter to editor). The dataset was then split randomly into a training-set (20% of total dataset) and test-set (n=4093)(Supplementary Figure.1). The final outcome was compared with manual decisions, revealing an accuracy of 80%. All studies included by MLA and/or human reviewer were subjected to full-text review by two different reviewers (HA & SB) to ensure eligibility. The machine learning was done using "sklearn" package using "LogisticRegression model" and the code is available at https://github.com/Hawra480/ML_updated1 and was run using python3.[2]

#### Data extraction and processing

We extracted the main biological and geographical moderators (e.g "country", "host type", "parasite species" "year of publication", "study type" and "study ID") using partially automated python scripting, followed by full manual review of extracted data to ensure accuracy. The extraction of prevalence data (extracted as "total sample size" and "positive cases") was manual extracted. Prevalence data was broken down to the taxa level of both the host and the parasite, with an average of 2-3 prevalence data extracted from each study. The exact host species was not available for all studies, thus, we conducted all relevant meta-regression models using the broad host categories (i.e. pigs, dogs, cats, camels, cattle, sheep, fish, raccoons, buffaloes, rodents, true bugs, flies, bees, birds, bats, fleas) and not the host-taxa level. Meta-regression models on bees, on the contrary, were conducted with the exact host taxa of bees-since it was available for all studies. Studies that report an overall prevalence data among several host categories without enough information to further break it down to each host level, were grouped under "animals". This group also includes host taxa found in single studies (such as frogs) as meta-regression models would otherwise be meaningless for that taxa. Sample size was standardised to the number of "individuals" sampled. Therefore, prevalence data among flies and bees, which are usually reported either as number of "pools" and number of "colonies" receptively, was estimated to "individual" number based on information reported to reflect the reported prevalence in each study. A phylogenetic representation of host taxa was created using "rotl" and "ggtreeExtra" packages[3–5] in R.[6]

#### Source data for qualitative and quantitative assessments in the main manuscript and supplementary figures

met-regression A (Table-1), met-regression B(Figure 3A), meta-regression C (figure 3B), meta-regression G (supplementary Figure 8) and the qualitative assessment for the geographic burden of trypanosomatid infection (supplementary Figure 3 and 5) were all conducted using the full data-set (n=581, K=1025).[7–585] The remaining meta-regressions were conducted on subgroups: meta-regression D (n=581, K=1025),[22,40,43,51,58,60,64,69,91,99,105–107,119,132,134,150,153,172,177,197,198,201,221,250–254,256,257,262,263,265,272,273] meta-regression E,[9–11,19,26,27,37,46,53,54, 72,76,83,89,93, 94,97,100,103,108,118,125,128,129,137–140,146,149,152–154,158,159,171,176,179,180,187,189,192,205,206,208,213,217,218,221, 231,235,246,248,249,255–257,260,261,267–270, 290,298,301,309,317,320,324, 327,334,344,347,349,361,368,393,414,422,425,429,444,446, 456,467,476,477,483,487,498, 500,508,514,517,529,533,539,547,548,568,584,586]

meta-regressions F1,[10,11,19,22,26,27,37,40,43,46,51,53,54,58,60,64,69,72,76,83,89,91,93,94,97,99,100,103,105–108,118,119,125,128,129,132,134,137–140,146,149,150,152–154,158,159,171,172,176,177,179,180,187,189,192,197,198,201,205,206,208,213,217,218,221,231,235,246,248–257,260–263,265,267–270,272, 273,290,298,301,309,317,320,324,327,334,344,347,349,361,368,393,414,422,425,429,444,446,456,467,476,477,483,487,498,500,508, 514, 517,529,533,539,547,548,568,584,586]

meta-regression F2,[7,8,12–18,20,21,23–26,28,29,31–36,38,39,41,42,44,45,47–50,52,55–57,59,61–63,65–68,70,71,74,75,77–82,84–88,90,92,95,96,98,100–102,104,109–117,120–127,130,131,133,135,137,141–145,147,148,151,152,155,156,160–170,173–176,178,181–186,188,190,191,193–196,199,200,202–204,207,209–216,219,220,222–230,232–234,236–240,242–245,247,259,266,271,275–289,291–297,299,300,302–319,321–323,325,326,328–332,335–342,345,346,348,350–360,362–367,369–392,394–413,415–421,423–428,430–443,445,447–455,457–466,468–475,478–486,488–497,499,501–506,508–513,515–528,530–532,534–538,540–546,548–562,564,566,567,569–583,585,587]

meta-regression F3,[10,11,19,22,26,27,37,40,43,46,51,53,54,58,60,64,69,72,76,83,89,91,93,94,97,99,100,103,105–108,118,119,125,128,129,132,134,137–140,146,149,150,152–154,158,159,171,172,176,177,179,180,187,189,192,197,198,201,205,206,208,213,217,218,221,231,235,246,248–257,260–263,265,267–270,272,273, 290,298,301,309,317,320,324,327,334,344,347,349,361,368,393,414,422,425,429,444,446,456,467,476,477,483,487,498,500,508,514,517,529,533,539,547,548,568,584,586] meta-regression F4,[9–11,19,26,37,40,53,54,76,83,89,93,94,97,100,118,125,128,129,137,138,149,152,154,159, 176,179,180,187,189,192,205, 208,213,217,221, 231,235,246,248,249,256,260,261,267–269,290,298,309,334,347,361,368,422,425,429,446,456,467,477,483,487,498,500,514,529,533,539,547,586] meta-regression F5,[27,46,72,103,108,139,140,146,153,158,171,206,218,255–257,301,317,320,324,344,349,393,414,444,476,508,517,548,568,584] meta-regression F6,[8–16,18–21,23–27,29,31–33,36–39,41,42,44–50,52–56,59,61–63,65–68,70–72,74–90,92–98,100–102,104,108,110–118,120–123,125,126,128–131,133,135,137–149,151–156,158–171,173–176,178–182,184–196,199,200,202–221,223–240,242–249,255,259–261,266–271,275,276,278,279,281–291,293–295,297–301,303–422,424–489,491–562,564,566–587]

meta-regression F7,[10–12,14,18–20,25,26,29,31,36,37,44,45,47–49,53–55,61,62,65,66,68,70,71,78,79,83–86,88–90,92,93,97, 98,101,102,113,117,122,123,126,128,129,135,137,141–143,147,154,156,164,167–169,186,189–191,193,194,196,203–205,207,208,210,211,213,216,217,219,232,238,246,248,249,261,267,275,276,278,279,281,283–285,287,290,291,294,295,298–300,306–310,312,313,315,318,322,327–329,331–334,336,339,341,343,345,347,348,350,353,355,357–363,365–368,371–373,375,376, 378,379,383,386,387,389,390,392,395,398,400–402,404,409,410,413,417,422,425,429,430,432–435,438–441,443,445,446,448,449,452–455,457–459,462,463,465–469,472, 474,475,477,481,482,485,489,491,493,494,497–502,507,509–512,514,515,521,523,525,529,532,535–540,543,549,551,552,554,555,557–559,564,566,567,571,573–577,579,580,586,587]

meta-regression F8,[8,9,13,15,21,23,24,33,38,39,41,42,56,67,74–77,82,87,94–96,100,104,108,110–112,114,115,118,120,121,125,130,131,133,138,145,148,149,152,159–163,166,170,171,173–176,178–182,184,185,187,188,192,195,200,202,209,214,215,220,221,223–231,233–237,239,240,242–245,247,259,260,266,268,269,277 ,282,288,289,

[297,303–305,311,314,316,319,321,326,330,337,338,340,342,349,354,356,364,369,370,377,380,381,384,385,391,396,397,399,405–408,411,412,418–421,424,426–428,431,436,437,442,447,450,451,456,460,461,464,470,479,480,483,484,486–488,496,505,511–513,516,519,520,522,524,526,527,530,531,533–535,541,542,544–547,550,553,560,561,572,578,581–583,585] meta-regression F9, [16,27,32,36,46,50,52,59,63,71,72,80,81,108,116,139,140,144,146,151,153,155,156,158,165,171,199,206,212,218,255,270,271,286,293,301,317,318,320,323–325,335,344,346,351,352,354,374,382,388,393,394,403,414–416,444,471 ,473,476,478, 492,493,495,501,503–506,508,517,518,528,548,556,562,568–570,579,584] meta-regression F10,[22,43,51,58,60,64,91,99,105–107,119,132,134,150,172,177,197,198,201,250–254,262,263,265,272,273] meta-regression F11,[22,43,51,58,60,64,91,99,105–107,119,132,134,150,172,177,197,198,201,250–254,262,263,265,272,273] and meta-regression F12.[22,43,51,58,60,64,91,99,105–107,119,132,134,150,172,177,197,198,201,250–254,262,263,265,272,273]

### Extended results

#### General model and publication bias

Out of 6046 unique citations, we have included 581 studies. Preforming the meta-analysis for all trypanosomatids infections (k=1025) reveals a low overall pooled estimate of (0.386, Cls: 0.367,0.405) back-transformed to a prevalence of (13.7%, Cls:12.4, 15.1), high heterogeneity rate (more than 90%) and no evidence of significant publication bias based on Egger's regression test for funnel plot asymmetry(p=0.726) with the predictor being the sample size as recommended for ecological and evolutionary studies with high heterogeneities (Supplementary Fig.6A-C). We also run separate meta-regression models with each of the sole moderators to assess the overall impact on the mean effect size. As expected, all tested moderators showed significant influence ($P<0.05$) on the mean effect size, as per the $Q_M$ statistics.

#### Geographic burden of trypanosomatids infections

The majority of the infections (60%) were reported from the American continents. In contrast, infections in Africa, Asia and Europe each contribute 12-13% of the total infections. The remaining 1% of quantitatively included data were from trypanosomatids infections in Australasia (Supplementary Fig.2A).

Considering the huge diversity of trypanosomatids species, it is perhaps not surprising that most countries exhibit a low range of an overall prevalence (1-20%)(Supplementary Fig.2B), with the exception of some geographic hot spots for trypanosomatids infections. An example of these hot spots include Chile and Ukraine with an average prevalence of 50-70%. This is followed by a 40-50% prevalence in Belgium, Germany and Tunisia and lastly between 20-40% in Algeria, Norway, Netherlands, United Kingdom and United States.

Interestingly, these geographic hot spots are usually caused by 1-2 groups of trypanosomatids species (i.e. either due to concentrated reporting and/or high transmission dynamics in an ecological niche). For instance, high prevalences in Algeria is mainly caused by *Leishmania* and *Trypanosoma evansi* infections (Supplementary Fig.5A & Fig. 5D). Similarly, high infection rates in Chile is mainly affected by honeybee parasites (i.e. *C. mellificae* and *L. passim*) and *Trypanosoma cruzi*(Supplementary Fig.5K & Supplementary Fig. 5B). On the other hand, prevalence in India seems to be predominantly influenced by high prevalence of multiple Trypanosomatidae-groups: *Leishmania* species, *Trypanosoma evansi* and bee parasites (i.e. *C. mellificae*, *L. passim*, *C. bombi* and *C. expoeki*) (Fig.5A & Fig. 5D & Fig. 5K & Fig. 5L ).

#### Supplementary Figures

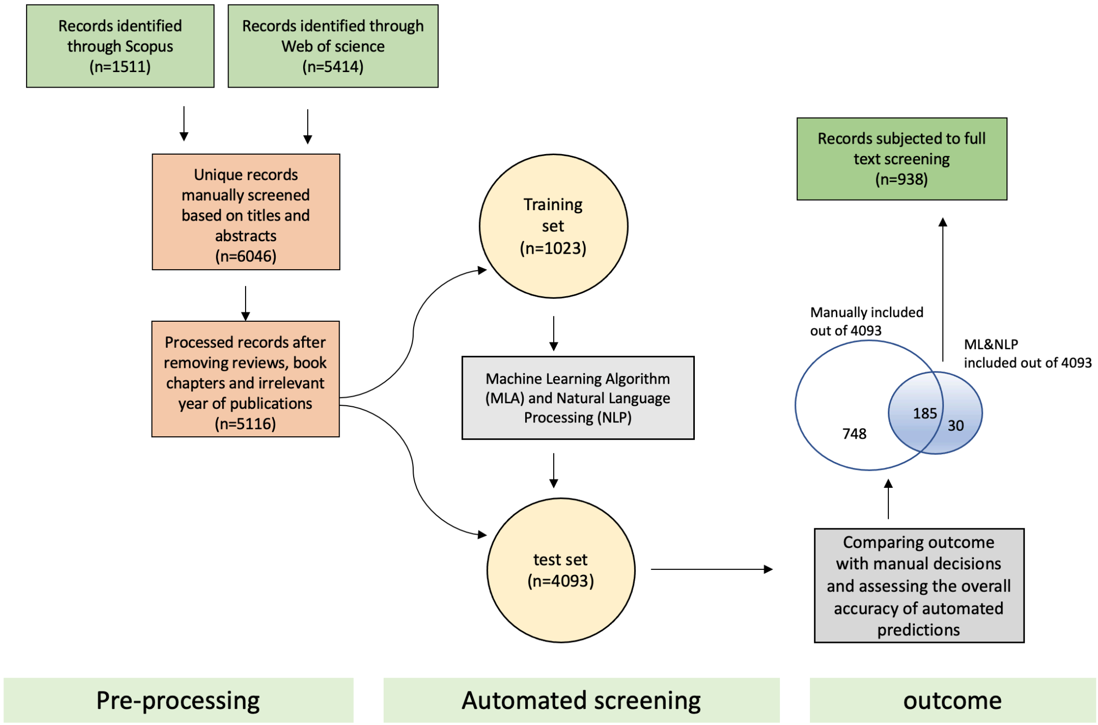


***Supplementary Figure 1:*** *Automated screening protocol.*

Summary of the main steps towards automated screening protocol as has been utilised in this study.

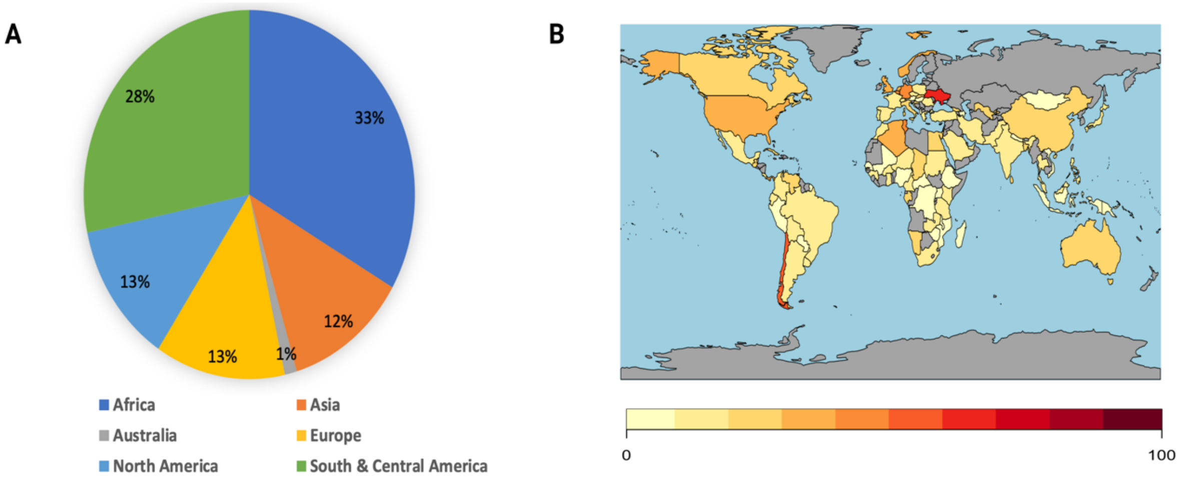


***Supplementary Figure 2:*** *A summary of spatial data .*

Sections A&B represent spatial burden of all included data (k=1025). With a geographic distribution of included data in different continents shown in section A, a world-map representation of all Trypanosomatid infections shown in section B.

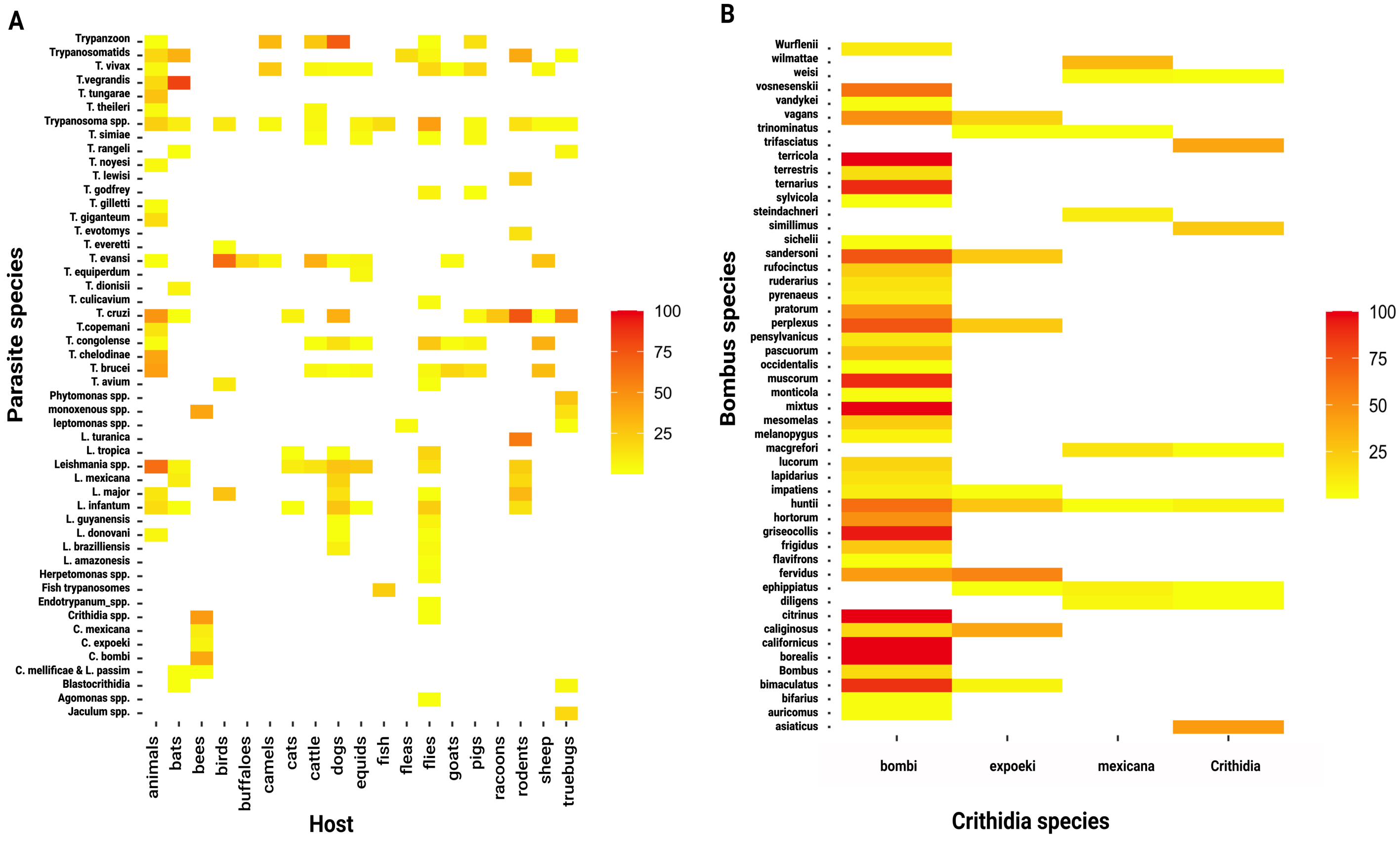


***Supplementary Figure 3:*** *A summary of prevalence data among various systems.*

A heat-map of the average prevalence among each host-parasite system is shown in (A), and a heat-map of the average prevalence among each of bumblebee-trypanosomatids system is shown in (B).

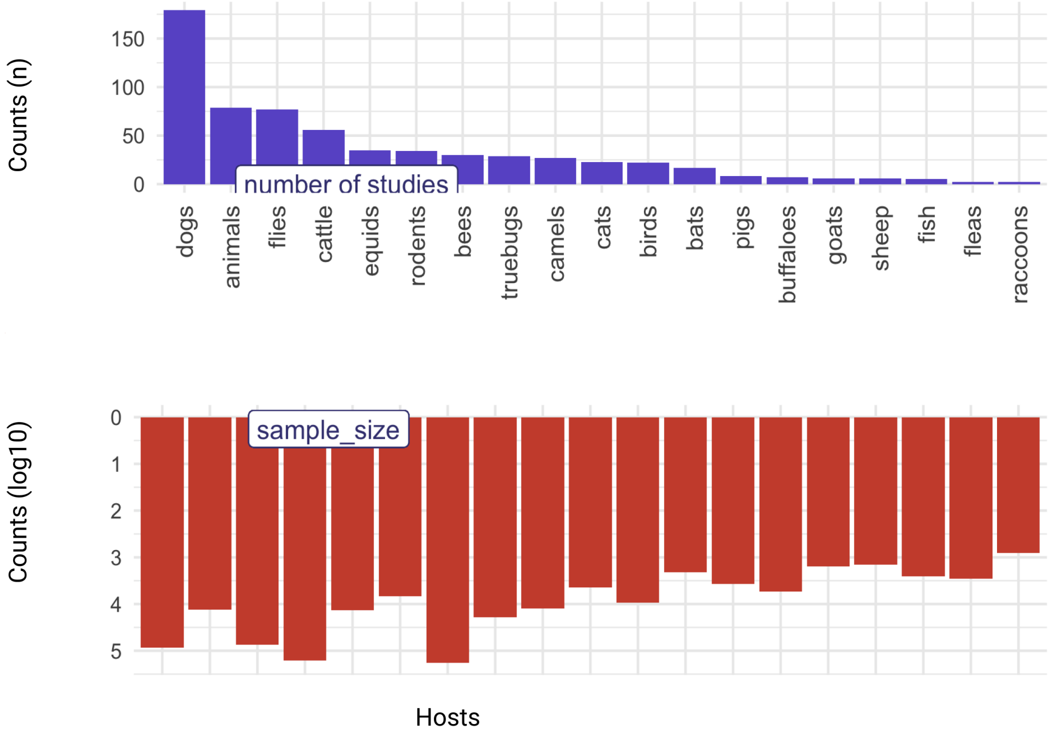


***Supplementary Figure 4:*** *A summary of host data*

Sections A& B outline temporal and numerical summary of all quantitatively included data (k=1025).a numerical break-down of aggregated data (k=1025) from 2000-2020 in section A and number of studies and total sample size among host groups shown in section B.

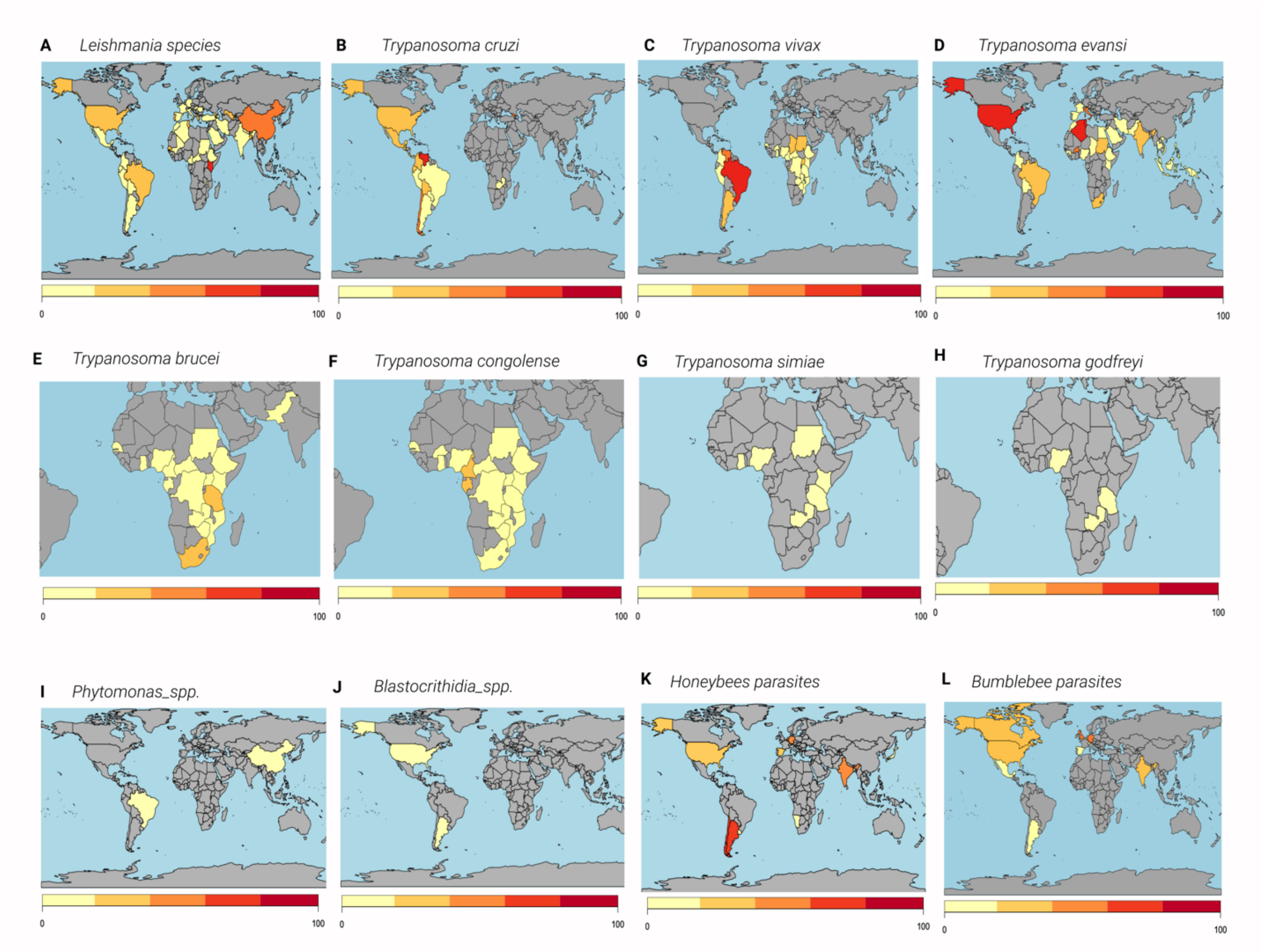


***Supplementary Figure 5:*** *Geographic distribution of various Trypanosomtidea-groups*

Sections (A-D) highlight the geographic burden of major dixenous-groups, sections (E-H) are the main dixenous-species usually constrained to African continent, and sections (I-L) highlight disease burden of neglected or rare Trypanosomtidea-groups. 5

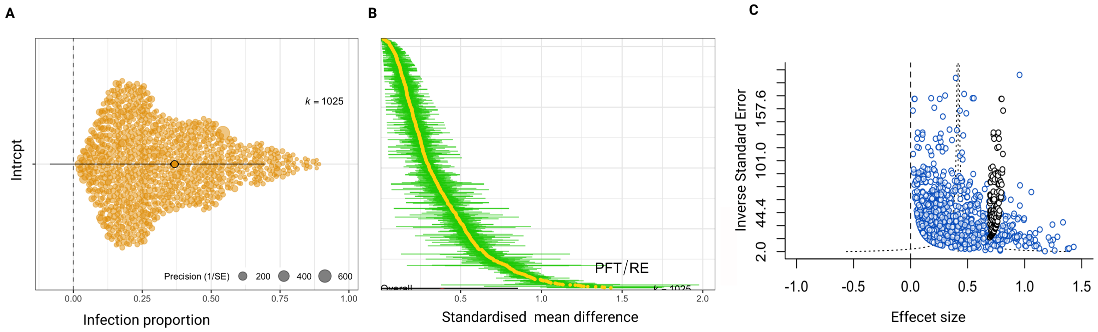


***Supplementary Figure 6:****: Forest and funnel plots of the general model*

A) A forest plot showing all quantitatively included evidence (k= 1023) with an overall pooled estimate of 0.333 (95% CI: 0.309, 0.359); back-transformed to a prevalence of 10.17%. (B) A visual representation of within and between study heterogeneity of all included evidence,(C) funnel plot with all included data shown as blue dots (k=1023) and estimated missing data shown as white dots.

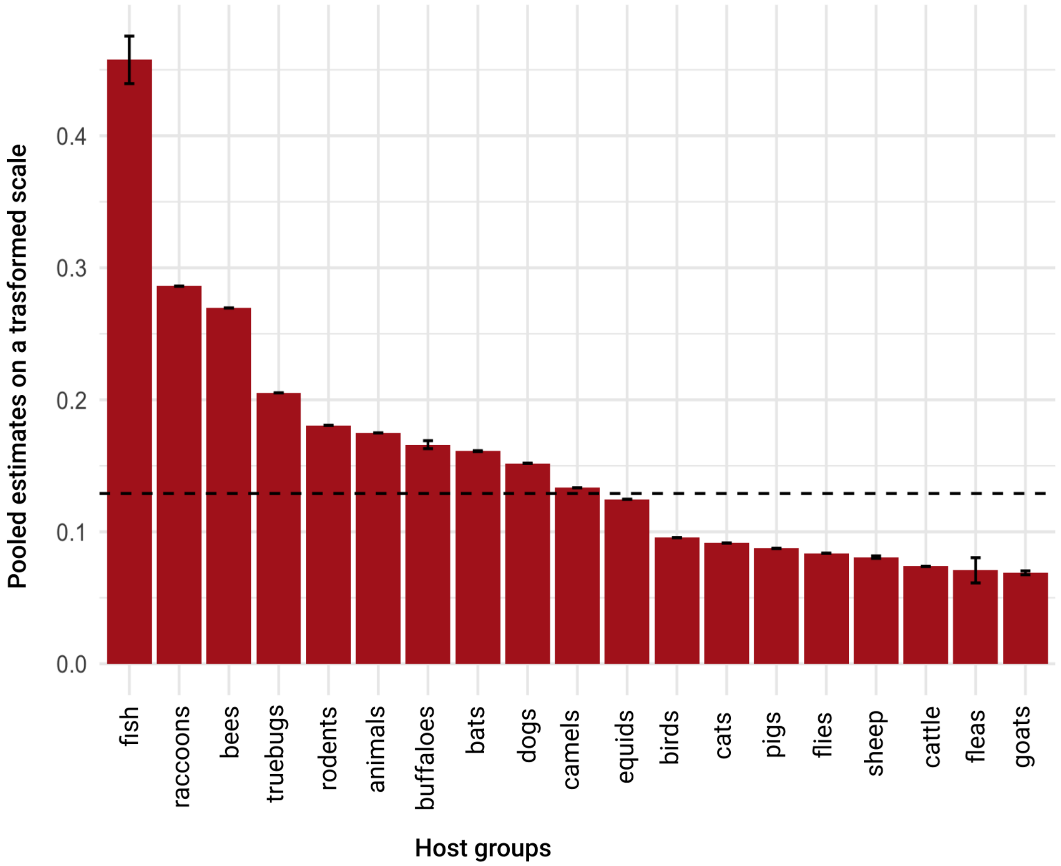


***Supplementary Figure 7:*** *Pooled proportion of infection in various host groups*

Pooled prevalence among each host group. Animals, as a group, includes all animals not included in the other groups and mixed groups of animals with no enough information to break them down to the host level.
