## Supplementary Table for "Double trouble: having two hosts reduces infection prevalence in vectored trypanosomatids"

**Supplementary Table1**: ***Searching terms utilised in this study***

| **Searching terms for all databases in (Web of Science)** | | **Hits** |
| --- | --- | --- |
| #1 | TS=(diversity OR incidence rate OR prevalence OR epidemiology)  Indexes=SCI-EXPANDED, SSCI, A&HCI, CPCI-S, CPCI-SSH, BKCI-S, BKCI-SSH, ESCI, CCR-EXPANDED, IC Timespan=All years | 1,811,220 |
| #2 | TS=(Trypanosomatids OR Trypanosoma)  Indexes=SCI-EXPANDED, SSCI, A&HCI, CPCI-S, CPCI-SSH, BKCI-S, BKCI-SSH, ESCI, CCR-EXPANDED, IC Timespan=All years | 36,226 |
| #3 | #2 AND #1  Indexes=SCI-EXPANDED, SSCI, A&HCI, CPCI-S, CPCI-SSH, BKCI-S, BKCI-SSH, ESCI, CCR-EXPANDED, IC Timespan=All years | 3,143 |
| #4 | TS= (Para-leishmania OR Leishmania OR Leptomonas OR Lotmaria OR Zelonia OR Borovskyia OR Crithidia OR Blastocrithidia OR Herpetomonas OR Phytomonas OR Wallaceina OR Jaenimonas OR Sergeia OR Paratrypansoma OR Lafontella OR Kentomonas OR Strigomonas OR Angomonas OR Blechomonas)  Indexes=SCI-EXPANDED, SSCI, A&HCI, CPCI-S, CPCI-SSH, BKCI-S, BKCI-SSH, ESCI, CCR-EXPANDED, IC Timespan=All years | 31,431 |
| #5 | #4 AND #1  Indexes=SCI-EXPANDED, SSCI, A&HCI, CPCI-S, CPCI-SSH, BKCI-S, BKCI-SSH, ESCI, CCR-EXPANDED, IC Timespan=All years | 2,656 |
| #6 combined search | #5 OR #3  Indexes=SCI-EXPANDED, SSCI, A&HCI, CPCI-S, CPCI-SSH, BKCI-S, BKCI-SSH, ESCI, CCR-EXPANDED, IC Timespan=All years | 5,414 |
| search stopped Jan 2020 | | |
| **Searching terms for database ( Scopus)** | | **Hits** |
| #1 | (diversity OR incidence rate OR prevalence OR epidemiology) AND (Trypanosomatids OR Trypanosoma) | 620 |
| #2 | ( ( diversity OR incidence AND rate OR prevalence OR epidemiology ) AND ( para-leishmania OR leishmania OR leptomonas OR lotmaria OR zelonia OR borovskyia OR crithidia OR blastocrithidia OR herpetomonas OR phytomonas) ) | 777 |
| #3 | (Wallaceina OR Jaenimonas OR Sergeia OR Paratrypansoma OR Lafontella OR Kentomonas OR Strigomonas OR Angomonas OR Blechomonas) | 152 |
| #4 combined search | #1 OR #2 OR #3 | 1511 |
| search stopped Jan 2020 | | |

**Supplementary Table 2: *Summary of the meta-regressions***

| Regression | Moderator | Outer random | Inner random | Intercept | Description | presentation | Data source |
| --- | --- | --- | --- | --- | --- | --- | --- |
| A | Parasite group. | Study ID | Host type, parasite type | No | Summary purpose | Table1 | ***Full dataset(581 studies)*** |
| B | Life-history of the parasite (dixenous, monoxenous, mixed). | Study ID | Host type, parasite type | Yes | Test whether monoxenous and dixenous parasites have different infection prevalence. | Fig3A | ***Full dataset(581studies)*** |
| C | Host group (insect, non-insect). | Study ID | Host type, parasite type | yes | Test whether insect hosts are more or less commonly infected than non-insect hosts of dixenous trypanosomatids. | Fig3B | ***Full dataset(581studies)*** |
| D | Host genus (insects infected with monoxenous trypanosomatids only). | Study ID | Host type, parasite type | yes | To compare insect hosts for differences in infection prevalence. | Fig4A | ***On subgroup of the dataset (36 studies)*** |
| E | Host genus (insects infected with dixenous trypanosomatids only). | Study ID | Host type, parasite type | yes | To compare insect hosts for differences in infection prevalence. | Fig4B | ***On subgroup of the dataset(102 studies)*** |
| F1 | Diagnostic method (insects only). | Study ID | Host type, parasite type | yes | Does diagnostic method affect the recorded prevalence. Because insects and non-insects use different methods, we separated these analyses. | Table2 | ***On subgroup of the dataset(134 studies)*** |
| F2 | Diagnostic method (non-insects only). | Study ID | Host type, parasite type | yes | Does diagnostic method affect the recorded prevalence in non-insect hosts. | Table2 | ***On subgroup of the dataset(448 studies)*** |
| F3 | Life-history of the parasite (dixenous, monoxenous) in insects only. | Study ID | Host type, parasite type | yes | Do monoxenous and dixenous trypanosomatids have different infection prevalence in insects. | Table2 | ***On subgroup of the dataset(134studies)*** |
| F4 | Life-history of the parasite (dixenous, monoxenous) in flies only. | Study ID | Host type, parasite type | yes | Do monoxenous and dixenous trypanosomatids have different infection prevalence in flies. | Table2 | ***On subgroup of the dataset(71 studies)*** |
| F5 | Life-history of the parasite (dixenous, monoxenous) in true bugs. | Study ID | Host type, parasite type | yes | Do monoxenous and dixenous trypanosomatids have different infection prevalence in true bugs. | Table2 | ***On subgroup of the dataset(32 studies)*** |
| F6 | Host-group (insects and non-insects) in dixenous trypanosomatids only | Study ID | parasite type | yes | Do dixenous trypanosomatids have different infection prevalence in insects. | Table2 | ***On subgroup of the dataset(520 studies)*** |
| F7 | Host-group (insects and non-insects) in Leishmania only. | Study ID | parasite type | yes | Does Leishmania have different infection prevalence in insects. | Table2 | ***On subgroup of the dataset(252 studies)*** |
| F8 | Host-group (insects and non-insects) in Trypanosoma (excluding T. cruzi) only. | Study ID | parasite type | yes | Does Trypanosoma have different infection prevalence in insects | Table2 | ***On subgroup of the dataset(200studies)*** |
| F9 | Host-group (insects and non-insects) in Trypanosoma cruzi only. | Study ID | parasite type | yes | Does T. cruzi have different infection prevalence in insects. | Table2 | ***On subgroup of the dataset(82 studies)*** |
| F10 | Wild vs managed bees | Study ID | Bee taxa, parasite taxa | yes | Do wild and managed bees have different infection prevalence. | Table2 | ***On subgroup of the dataset (30 studies)*** |
| F11 | Wild vs managed bumblebees | Study ID | Bee taxa, parasite taxa | yes | Do wild and managed bumblebees have different infection prevalence. | Table2 | ***On subgroup of the dataset(11 studies)*** |
| F12 | Wild vs managed honeybees | Study ID | Bee taxa, parasite taxa | yes | Do wild and managed honeybees have different infection prevalence. | Table2 | ***On subgroup of the dataset(20 studies)*** |

**Supplementary Table 3: Corrected cut-off of significance**

| Description of null hypothesis | Meta-regression ID | p-value(k) | k | Adjusted cut-off after correction for multiple testing | conclusion |
| --- | --- | --- | --- | --- | --- |
| No significant difference between infections diagnosed via molecular versus (microscopic & culture based) tools among non-insects. | F2.1 | <.0001 | 1 | 0.002 | sig |
| No significant difference in infections diagnosed via serological versus (microscopic & culture based) tools among non-insects. | F2.2 | <.0001 | 2 | 0.004 | sig |
| No significant difference in *Leishmania* infections among insects and non-insects. | F7 | 0.0001 | 3 | 0.006 | sig |
| No significant difference in infections diagnosed via molecular versus (microscopic & culture based) tools among insects. | F1 | 0.0013 | 4 | 0.008 | sig |
| No significant difference between ‘monoxenous’ infections and ‘dixenous’ infections among all hosts. | B.1 | 0.0057 | 5 | 0.01 | sig |
| No significant difference in *dixenous* infections among insects and non-insects. | F6 | 0.0067 | 6 | 0.012 | sig |
| No significant difference between Dixenous infections and Monoxenous infections among insects. | F3 | 0.0067 | 7 | 0.014 | sig |
| No significant difference in infections between insects and non-insects. | C | 0.067 | 8 | 0.016 | Not sig |
| No significant difference in infections between Trypanosoma and Endotrypanum genera | E.2 | 0.0727 | 9 | 0.018 | Not sig |
| No significant difference between ‘mixed trypanosomatids’ infections and ‘dixenous’ infections among all hosts. | B.2 | 0.0889 | 10 | 0.02 | Not sig |
| No significant difference in infections caused by Blastocrithidia_spp.vs ‘jaculum’_spp. | D.2 | 0.1865 | 11 | 0.022 | Not sig |
| No significant difference in prevalence of infections reported from insects and non-insects caused by *Trypanosoma spp. (*excluding *T.cruzi)* | F8 | 0.241 | 12 | 0.024 | Not sig |
| No significant difference in infections caused by Agomonas_spp. Vs ‘jaculum’_spp. | D.1 | 0.2714 | 13 | 0.026 | Not sig |
| No significant difference in infections caused by Leptomonas_spp. Vs ‘jaculum’_spp. | D.5 | 0.2829 | 14 | 0.028 | Not sig |
| No significant difference in infections caused by Endotrypanum or leishmania genera. | E.1 | 0.2937 | 15 | 0.03 | Not sig |
| No significant difference in *T.cruzi’s* infections among insects and non-insects | F9 | 0.3396 | 16 | 0.032 | Not sig |
| No significant difference in infections caused by Herpetomonas_spp. Vs ‘jaculum’_spp. | D.4 | 0.3654 | 17 | 0.034 | Not sig |
| No significant difference in infection prevalence caused by dixenous *spp.* compared to *m*onoxenous *spp among flies.* | F4 | 0.4566 | 18 | 0.036 | Not sig |
| No significant difference in infections between Phytomonas and Endotrypanum genera | E.3 | 0.4869 | 19 | 0.038 | Not sig |
| No significant difference in infections among managed and wild bees. | F10 | 0.7268 | 20 | 0.04 | Not sig |
| No significant difference in infection **between dixenous *spp.*and monoxenous** *spp among true bugs.* | F5 | 0.7883 | 21 | 0.042 | Not sig |
| No significant difference in infections among managed and wild honeybee. | F11 | 0.8357 | 22 | 0.044 | Not sig |
| No significant difference in infections caused by Crithidia_spp. And ‘jaculum’_spp. | D.3 | 0.8438 | 23 | 0.046 | Not sig |
| No significant difference in infections among managed and wild bumblebees. | F12 | 0.9096 | 24 | 0.048 | Not sig |
| No significant difference in infections caused by monoxenous vs ‘jaculum’_spp. | D.6 | 0.9372 | 25 | 0.05 | Not sig |

Number of comparisons from all meta-regressions(m=25), adjusted cut-off based on Benjamini-Hochberg method, the formula is (k/m)*0.05.

Where K is the index of p-value(K), arranged from smallest to largest, and m is the number of comparisons. We concluded that there is a significant p-value when p-value(k)< Adjusted cut-off. Whenever this is not met (p-value(k)> Adjusted cut-off) we conclude non- significance for all bigger p-values(k).

D.1-D.7 represents statistical comparisons of each level with the intercept level within a moderator within meta-regression D. Same goes for F1.1-F1.2 (meta-regression F1), E.1 - E.2 (meta-regression E) and B.1 and B.2 (meta-regression B).
